# Mitorubin, berberrubine-based compounds that improve mitochondrial function, exhibit cardioprotective effects against age-related cardiac dysfunction

**DOI:** 10.1101/2025.05.01.651794

**Authors:** Michio Sato, Daishi Tanabu, Daisuke Torigoe, Tsuyoshi Kadomatsu, Keito Taniwaka, Yoshionbu Ogata, Isshin Shiiba, Yuiko Suzuki, Ryoko Inatome, Takeshi Tokuyama, Toshihiko Takeiwa, Satoshi Inoue, Eito Kanai, Takashi Hamano, Hiromi Hirata, Kayoko Kanamitsu, Hiroyuki Kusuhara, Akihito Yokosuka, Yoshihiro Mimaki, Hideki Abe, Yuichi Oike, Shigeru Yanagi

## Abstract

Mitochondria play a central role in cellular energy metabolism and homeostasis, and their dysfunction is closely linked to the progression of age-related diseases. The mitochondrial ubiquitin ligase MITOL (also known as MARCHF5) is a key regulator of mitochondrial dynamics and function, and reduced MITOL expression in the mouse heart has been implicated in mitochondrial dysfunction and cardiac aging. In this study, we identified berberrubine as a compound that promotes MITOL expression and activates mitochondria. We further assembled a group of berberrubine-based compounds, including its quinoid form and a newly developed water-soluble derivative, and collectively named them “Mitorubin” as mitochondria-activating compounds with therapeutic potential. While conventional berberrubine has poor water solubility, the addition of acetic acid significantly improved its solubility, enabling formulation as a solution. Mitorubin enhanced MITOL expression in cultured cells, increased mitochondrial DNA content and expression of mitochondrial proteins, and promoted mitochondrial respiration. In a model of age-related cardiac dysfunction, oral administration of Mitorubin restored mitochondrial function, improved cardiac performance, suppressed myocardial hypertrophy, and alleviated pulmonary congestion. Moreover, Mitorubin did not shorten lifespan in aged mice and significantly extended lifespan in high-fat diet-fed mice, suggesting both safety and efficacy under chronic administration. These findings suggest that Mitorubin is a promising mitochondrial activator and may represent a novel therapeutic strategy for age-related diseases.

## Introduction

Mitochondria are essential organelles in eukaryotic cells that play a crucial role in various cellular functions, including energy metabolism, calcium homeostasis, lipid metabolism, and apoptosis. Structurally, mitochondria exhibit elongated, branched, or granular forms within cells, and their morphology is dynamically regulated by fusion factors such as Mfn1/2 and Opa1, as well as fission factors like Drp1, in response to changes in the cellular environment^1,2^. These morphological changes occur due to factors such as nutrient availability and mitochondrial damage^3,4^. Notably, in cardiomyocytes, aging is associated with an increased prevalence of fragmented mitochondria, suggesting a potential link between mitochondrial dynamics and age-related cardiac dysfunction^5,6^.

Mitochondria play a central role in ATP production through oxidative phosphorylation (OXPHOS), a process that primarily generates cellular energy while also producing reactive oxygen species (ROS), such as superoxide and hydrogen peroxide, as inevitable byproducts^7,8^. The excessive accumulation of ROS damages mitochondrial DNA (mtDNA) and proteins, contributing to aging and the progression of age-related diseases^9–11^. However, moderate levels of ROS play a physiological role in maintaining cellular homeostasis and promoting adaptive responses that optimize energy metabolism. This concept, known as mitochondrial hormesis (mitohormesis), suggests that mild mitochondrial stress can induce antioxidant responses and adaptive mitochondrial changes, ultimately enhancing cellular function^12,13^.

MITOL (also known as MARCHF5) is an E3 ubiquitin ligase that spans the mitochondrial outer membrane four times and regulates mitochondrial homeostasis by targeting proteins such as Drp1. MITOL suppresses excessive mitochondrial fission by promoting Drp1 degradation and thereby regulates mitochondrial dynamics^14^. Additionally, MITOL plays a broader role in cellular quality control by degrading pathogenic proteins that are involved in neurodegenerative diseases, such as mutant superoxide dismutase 1 (SOD1) and polyglutamine aggregates, as well as the microtubule-stabilizing factor MAP1B^15–17^. In cultured cells, MITOL deficiency leads to increased mitochondrial ROS production and premature cellular senescence. This aging phenotype has also been observed *in vivo*. Studies using cardiomyocyte-specific MITOL-knockout mice have revealed that MITOL downregulation disrupts mitochondrial dynamics through Drp1 accumulation, leading to cardiac aging^18^. Similarly, neuron-specific MITOL deficiency has been shown to exacerbate the pathology of Alzheimer’s and Parkinson’s diseases^19,20^. Given that MITOL expression declines significantly with aging, it has been proposed that reduced MITOL levels contribute to aging and age-related diseases, making MITOL a potential therapeutic target. However, the mechanisms that regulate MITOL expression *in vivo* remain largely unknown, and the identification of MITOL activators or inducers has been a major research challenge.

Age-related cardiac dysfunction is a condition characterized by structural and functional deterioration of the myocardium with aging, in which mitochondrial dysfunction plays a central role^21^. Mitochondria serve as the primary source of energy in cardiomyocytes, generating ATP through OXPHOS. However, aging alters mitochondrial morphology, dynamics, and function, leading to a decline in energy production capacity^10^. Specifically, disruptions in the balance between mitochondrial fusion and fission, impairment of the electron transport chain, and abnormal ROS accumulation contribute to cardiomyocyte damage^22^. Furthermore, age-related mtDNA damage and the decline of mitochondrial quality control mechanisms, such as autophagy and mitophagy, exacerbate heart failure progression^23,24^. In recent years, therapeutic strategies aimed at preserving and restoring mitochondrial function have gained increasing attention, prompting ongoing efforts to develop novel treatment approaches^25,26^.

Berberine is an isoquinoline alkaloid and a major bioactive component of goldthread (e.g., *Coptis japonica*) and Amur cork tree (e.g., *Phellodendron amurense*), which have long been used in traditional East Asian medicine as anti-diarrheal agents^27^. Berberine is known to exhibit diverse pharmacological activities, including antibacterial, anti-inflammatory, hypolipidemic, antihypertensive, and hypoglycemic effects^28^. Numerous studies have identified molecular targets of berberine, revealing its involvement in multiple cellular pathways. Interestingly, berberine has also been reported to influence mitochondrial function through mechanisms such as ROS suppression, regulation of mitochondrial dynamics, mitophagy activation, mitochondrial biogenesis, and calcium homeostasis^29^. These effects contribute to its pharmacological properties by maintaining mitochondrial quality and function.

In this study, we conducted an exploratory evaluation of a panel of compounds, including herbal extracts, and found that *Coptis japonica* and *Phellodendron amurens*e act as potent inducers of MITOL expression. Further analysis identified berberrubine, a primary *in vivo* metabolite of berberine, as a key regulator of mitochondrial protein expression and mitochondrial biogenesis. Berberrubine was also found to promote mitochondrial dynamics, increasing the proportion of elongated mitochondria, and significantly enhance mitochondrial respiration. Notably, berberrubine exhibits extremely poor water solubility, which limits its formulation versatility and oral usability. To overcome this challenge, we successfully synthesized a novel berberrubine acetic acid adduct with enhanced water solubility and demonstrated its efficacy in improving age-related cardiac dysfunction. Our findings suggest that we have identified a class of berberrubine-based compounds with mitochondrial-activating properties, which we designate as “Mitorubin”, a potential therapeutic approach for age-related diseases associated with mitochondrial dysfunction.

In the following sections, the individual components of Mitorubin are referred to by their solid-state structure-based names (quinoid-type berberrubine and berberrubine acetic acid adduct) or their corresponding abbreviations (Q-BRB and AcA-BRB) for clarity.

## Results

### 1. Identification of berberrubine, a berberine metabolite, as an inducer of MITOL expression

In our previous study, we demonstrated that MITOL expression in the mouse brain declines sharply during the postnatal period and then gradually decreases with aging^30^, suggesting its potential role as an age-related regulatory factor. To investigate whether this trend extends to the heart, we examined MITOL protein levels in mouse cardiac tissue and found a gradual age-dependent decline (Fig. 1a, b). This finding suggests that MITOL functions as an age-related factor in the heart as well.

**Figure 1.**
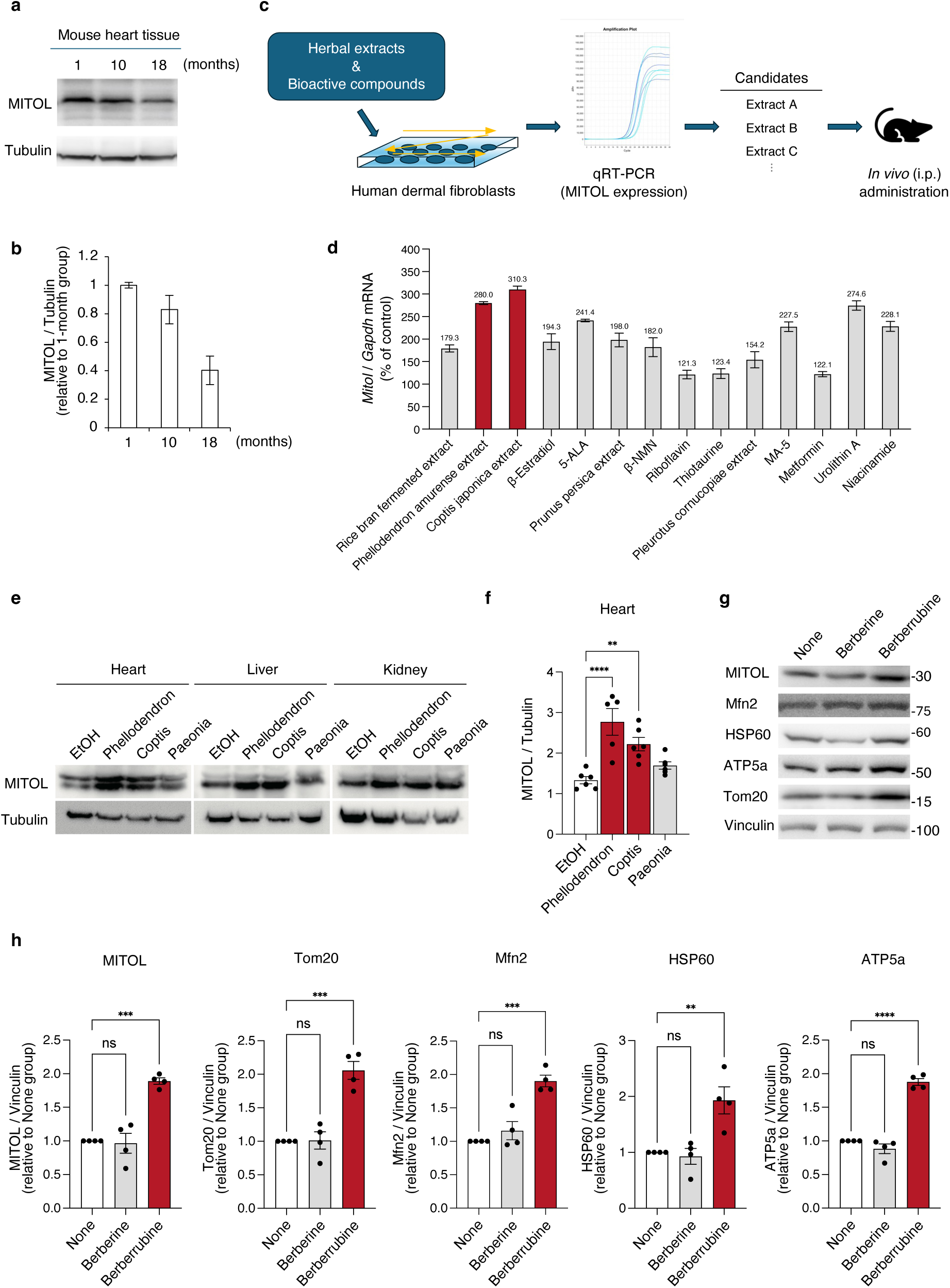
Identification of berberrubine, a berberine metabolite, as a compound that enhances MITOL expression. (**a, b)** Age-related decrease in MITOL expression. Western blot analysis of MITOL expression in the heart at each indicated month of age. Signal intensities were normalized to Tubulin and presented as values relative to the 1-month group. **(c)** Schematic representation of the compound evaluation strategy. Various herbal extracts and previously reported mitochondria-activating bioactive compounds were individually administered to human dermal fibroblasts. Candidate compounds identified based on MITOL mRNA upregulation by qRT-PCR were subsequently administered to mice to assess MITOL expression in multiple organs. **(d)** Relative MITOL mRNA levels in human dermal fibroblasts after 24-hour treatment with each compound, quantified by qRT-PCR. mRNA levels were normalized to GAPDH and expressed as a percentage relative to the control group (without test substances). Data represent the average of three independent experiments (n = 3). **(e, f)** Increased MITOL expression in mouse tissues following herbal extract administration. Mice were administered each herbal extract (*Phellodendron amurense*, *Coptis japonica*, or *Paeonia suffruticosa*) intraperitoneally, and 9 days later, MITOL expression levels in various organs were assessed by Western blot. **f** shows the graph of MITOL expression levels in the heart, normalized to tubulin (n = 5– 6 per group). *Paeonia suffruticosa* extract was included based on its prior effect on MITOL expression in hair follicle keratinocytes (data not shown); however, it did not increase MITOL expression in the heart, liver, or kidney in this experiment. **(g, h)** Upregulation of MITOL and mitochondrial proteins in C2C12 myoblasts following berberrubine treatment. C2C12 cells were treated with berberine (10 μM) or quinoid-type berberrubine (10 μM) for 48 hours, and protein expression levels were analyzed by Western blot (**g**). **h** shows the quantification of Western blot signals for MITOL, Tom20, Mfn2, HSP60, and ATP5a, normalized to Vinculin and expressed as values relative to the None group (n = 4 per group). The None group indicates cells cultured without any additives, including solvents. Statistical significance was determined using two-way ANOVA followed by Bonferroni post hoc test **(f)** or one-way ANOVA followed by Tukey’s HSD test **(h)**. \**p* < 0.05; \*\**p* < 0.01; \*\*\**p* < 0.001; \*\**p* < 0.0001; ns, not significant.

Next, to identify compounds capable of inducing MITOL expression, we evaluated a panel of herbal extracts and mitochondria-activating bioactive compounds for their effects on MITOL mRNA levels (Fig. 1c). Quantitative RT-PCR (qRT-PCR) analysis of human dermal fibroblasts treated with each test substance revealed that *Coptis japonica* and *Phellodendron amurense* extracts induced MITOL mRNA expression by approximately threefold (Fig. 1d). When these extracts were administered to mice, we observed increased MITOL protein expression across multiple organs, including the heart, in both *Coptis japonica* and *Phellodendron amurense* extract-treated groups (Fig. 1e, f).

Given that berberine is the major bioactive component of these extracts, we hypothesized that berberine itself might enhance MITOL expression. However, when C2C12 myoblasts were treated with berberine, MITOL expression did not increase. Instead, its major *in vivo* metabolite berberrubine^31^, which was used as the quinoid-type form (defined later in Fig. 2), significantly upregulated MITOL expression (Fig. 1g). Notably, berberrubine also markedly increased the expression of other mitochondrial-associated proteins (Fig. 1g, h), suggesting that berberrubine broadly influences mitochondrial protein regulation.

**Figure 2.**
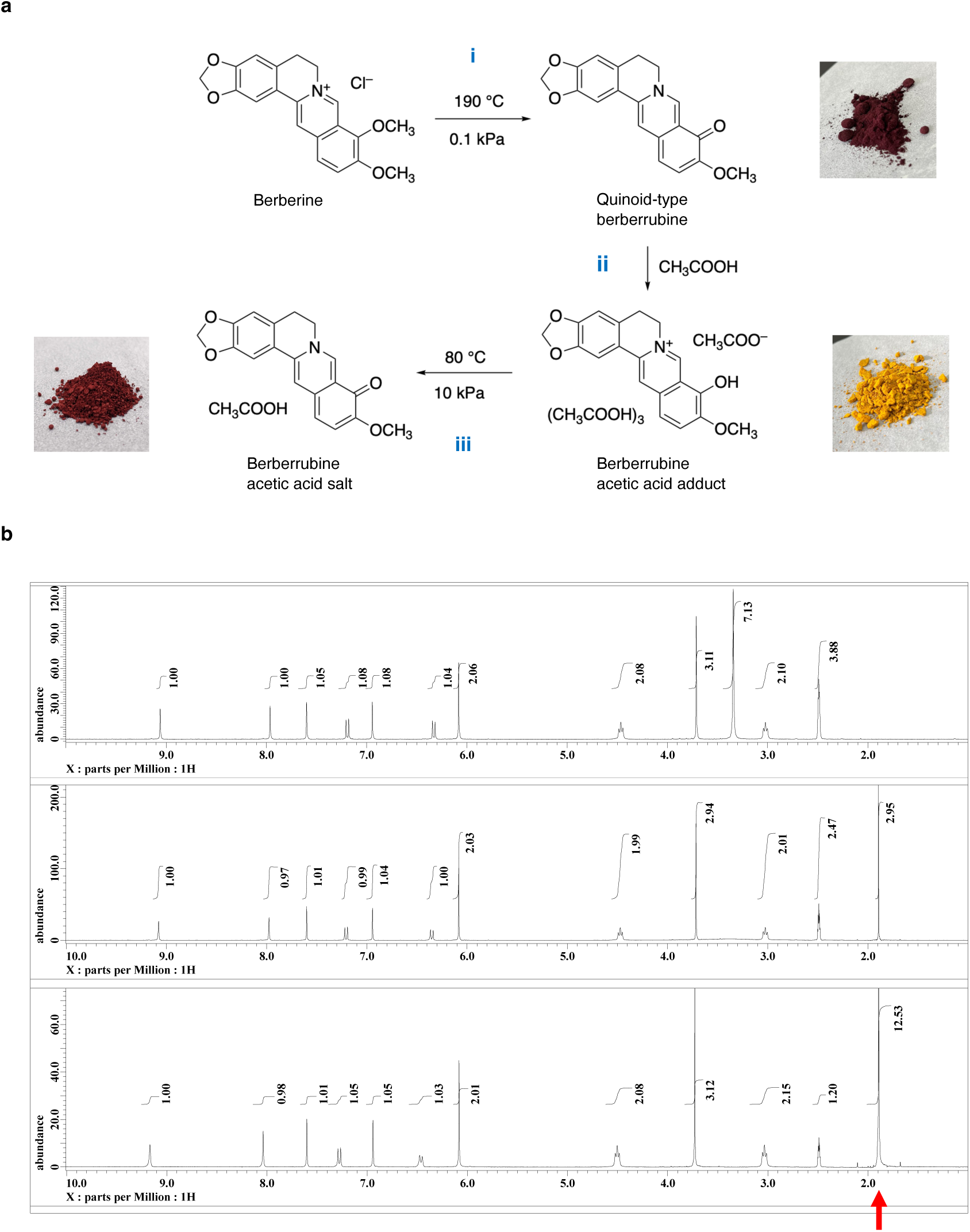
Synthesis of quinoid-type berberrubine, berberrubine acetic acid adduct, and berberrubine acetic acid salt. **(a)** Quinoid-type berberrubine, shown as a reddish-black solid, was obtained by thermal decomposition of berberine chloride under reduced pressure (Step i). The resulting berberrubine powder was suspended in acetic acid, and the precipitate formed was collected by suction filtration, washed with diethyl ether, and dried under reduced pressure (Step ii), yielding the yellow solid berberrubine acetic acid adduct. Further drying (Step iii) produced the reddish-brown solid berberrubine acetic acid salt. **(b)**¹H NMR spectra of the berberine-derived compounds in deuterated DMSO. From top to bottom: reddish-black solid (quinoid-type berberrubine), reddish-brown solid (berberrubine acetic acid salt), and yellow solid (berberrubine acetic acid adduct). The peaks indicated by the red arrow correspond to the protons of the methyl groups in acetic acid molecules.

### 2. Preparation of water-soluble berberrubine and its property

#### Water-soluble berberrubine: Preparation and structural analysis of berberrubine acetic acid adduct

The quinoid-type berberrubine derived from berberine (berberine chloride) is a water-insoluble compound^32^. To improve the biological value, especially to obtain water solubility of berberrubine, we produced numerous salts with various Brønsted acid. The quinoid-type berberrubine, which was used in the mitochondrial protein expression experiments described above, would be obtained by heating berberine chloride in DMF, then pouring the reaction solution into ice water and filtering the resulting solid. It is also possible to synthesize berberrubine in a short time by directly irradiating microwaves to berberine chloride using a microwave synthesis device. However, these synthesis methods make it difficult to produce berberrubine for biological studies in quantity. Therefore, we decided to synthesize it on a large scale via a method of pyrolysis by heating under reduced pressure (Fig. 2a, Step i). This method allowed the heating of up to about 200 g of berberine (berberine chloride) at a time in the laboratory, yielding quinoid-type berberrubine in quantitative amounts.

As noted above, the obtained berberrubine did not dissolve, even in 1,000 mL of water. Next, various salts were produced using different Brønsted acids to improve the water solubility of berberrubine. Among the many salts produced, it was found that the one obtained using acetic acid was extremely soluble in water. The preparation was carried out as follows, and two types of salt were obtained as a result.

When acetic acid was added to the powdered berberrubine and stirred, the solution turned dark reddish-brown, and a yellow precipitate gradually formed. The suspension was filtered by suction filtration while being washed with diethyl ether, and dried under reduced pressure to obtain a yellow solid (Fig. 2a, Step ii). As described later, the ^1^H NMR data of the resulting yellow solid in deuterated DMSO solution showed a spectrum with the exact same chemical shifts as quinoid-type berberrubine, as well as a spectrum containing about four molecules of acetic acid. In the NMR spectra of salts made from other Brønsted acids, the ratio of berberrubine to acids was almost 1:1, whereas the yellow solid made using acetic acid retained more acetic acid molecules. Therefore, the yellow crystals were dried under reduced pressure while being heated to further dry them, and a reddish-brown solid was obtained (Fig. 2a, Step iii). The ^1^H NMR spectrum of the reddish-brown solid was almost identical to that of the yellow solid, with the only difference being that the ratio of acetic acid in the reddish-brown solid was one molecule of acetic acid per one molecule of berberrubine.

#### Structural analysis of berberrubine acetic acid adduct and acetate salt

The ^1^H NMR spectra of the reddish-black berberrubine synthesized from berberine and the yellow and reddish-brown solids produced from berberrubine in deuterated DMSO solution differ only in the presence or absence of protons derived from acetic acid molecules and in the integral values, but the chemical shift values of the protons corresponding to berberrubine are the same (Fig. 2b). However, since the colors of the solids differ, it was presumed that berberrubine in each solid has a different chemical structure. Therefore, solid-state ^13^C NMR, which can directly observe the target nucleus in the solid state, was measured.

The ^13^C peak shape of the product obtained through pyrolysis, as measured by the CPMAS method, showed a ketone-derived C-13 peak (13), and was similar to the peak shape of the solution ^13^C-NMR spectrum reported in Figure 1 of^32^, so the product was presumed to be a quinoid-type berberrubine (Suppl. Fig. 1a).

The yellow solid was presumed to be phenol-type berberrubine because the ^13^C peak shape of the yellow solid measured by the CPMAS method was similar to that of the solution ^13^C-NMR spectrum of the phenol-type hydrochloride, and no ^13^C peaks derived from ketones were observed (Suppl. Fig. 1b). In addition, multiple peaks derived from acetic acid (25, 26) were observed, suggesting the presence of multiple acetic acids in different states. Of the four peaks (26), the high-frequency side (182 ppm side) was presumed to be derived from acetic acid with increased anionicity due to hydrogen bonds and salt formation, and the low-frequency side (172 ppm side) was presumed to be derived from acetic acid with low anionicity. These results indicate that the yellow solid is a phenol-type berberrubine, and furthermore that it is a solid containing a total of four molecules derived from acetic acid, namely one acetate ion and three acetic acid units. This result is consistent with the observation of protons derived from acetic acid equivalent to four molecules of acetic acid in DMSO-*d*_6_ solution NMR.

The ^13^C peak shape of the reddish-brown solid measured by CPMAS method was similar to that of quinoid-type berberrubine (Suppl. Fig. 1c) and the solution ^13^C-NMR spectrum reported in Figure 1 of^32^, The reddish-brown solid was therefore presumed to be a quinoid-type structure. In addition, several peaks were also observed that were derived from acetic acid (25, 20), but these were mainly at low frequencies (172 ppm) and were presumed to be derived from acetic acid, which possesses a low anionic property.

From these results, it was determined that the reddish-brown solid is a salt in which one molecule of acetic acid has acted on quinoid-type berberrubine (berberrubine acetate salt), and the yellow solid is a substance in which four molecules of acetic acid have acted on phenol-type berberrubine (berberrubine acetate adduct, Fig. 2a).

It was revealed that the formation of Brønsted acid salts of alkaloids, which are nitrogen-containing compounds, occurs through protonation or hydrogen bonding of the organic acid to the nitrogen atom, with the ratio of Brønsted acid to alkaloid being approximately 1:1, whereas the reaction of berberrubine with acetic acid results in the construction of a crystalline substance in a ratio of 1:4, and that drying these under reduced pressure produces normal Brønsted acid salts.

#### Enhanced water solubility of berberrubine through acetic acid modification

The solubility of three compounds, the reddish-black solid (quinoid-type berberrubine), the reddish-brown solid (berberrubine acetic acid salt), and the yellow solid (berberrubine acetic acid adduct), in water was roughly examined. The liquid NMR data of the three in deuterated DMSO solvent were very similar, but their water solubilities were significantly different. Because quinoid-type berberrubine possesses very high lipophilicity^32^, the reddish-black solid was not possible to observe any dissolution even in a very small amount in 1,000 mL of water. On the other hand, the reddish-brown solid dissolved in about 30 mg in 1 mL of water.

Surprisingly, the yellow solid showed a very high water-solubility, dissolving about 1 g in 1 mL of water. In this way, we succeeded in significantly improving the solubility of berberrubine in water by reacting quinoid berberrubine with acetic acid. This also makes it possible to conduct animal experiments using drinking water.

### 3. *In vitro* ADME evaluation of quinoid-type berberrubine and berberrubine acetic acid adduct

Next, to evaluate the physicochemical properties of quinoid-type berberrubine (Q-BRB) and berberrubine acetic acid adduct (AcA-BRB) synthesized from berberine chloride, we conducted an *in vitro* ADME evaluation (Table 1).

**Table 1.** *In vitro* ADME evaluation of quinoid-type berberrubine and berberrubine acetic acid adduct. Membrane permeability, liver microsomal metabolic stability, and plasma protein binding of quinoid-type berberrubine (Q-BRB) and berberrubine acetic acid adduct (AcA-BRB) were evaluated. In the membrane permeability section, the pH values of the donor and acceptor compartments are provided.

| Compounds | Membrane permeability<br>[$\times 10^{-6}$ cm/sec] | | | |
| --- | --- | --- | --- | --- |
|  | pH4.5/4.5 | pH6.4/7.4 | pH7.4/7.4 | pH8.6/8.6 |
| Q-BRB | — | 1.63 | 2.90 | 2.60 |
| AcA-BRB | 0.220 | 1.87 | 1.80 | — |

**Table 1.**978 ***In vitro* ADME evaluation of quinoid-type berberrubine and berberrubine acetic acid adduct.**
| Compounds | Liver microsomal metabolic stability<br>[mL/min/kg] |  | Plasma protein binding<br>(fraction unbound) |  |
| --- | --- | --- | --- | --- |
|  | human<br>(pH7.4) | mouse<br>(pH7.4) | human<br>(pH7.4) | mouse<br>(pH7.4) |
| Q-BRB | < 22.0 | 1839 | 0.359 | 0.162 |
| AcA-BRB | < 22.0 | 1794 | 0.374 | 0.203 |

#### Membrane permeability

Membrane permeability was assessed using the parallel artificial membrane permeability assay (PAMPA). The permeability of quinoid-type berberrubine at pH 8.6/8.6 (donor/acceptor) was comparable to that of metoprolol, a reference drug with approximately 95% human intestinal absorption. Both Q-BRB and AcA-BRB exhibited similar permeability under pH 6.4/7.4 and pH 7.4/7.4 conditions, comparable to Q-BRB at pH 8.6/8.6. These findings suggest that both compounds predominantly exist in the quinoid form at pH ≥ 6.4. In contrast, the permeability of AcA-BRB at pH 4.5/4.5 was markedly lower, which is consistent with a previous report^32^.

#### Metabolic stability

Metabolic stability was evaluated using liver microsomes from mice and humans. Both compounds exhibited low metabolic stability in mouse liver microsomes. In contrast, in human liver microsomes, both compounds showed markedly higher metabolic stability. Berberrubine is known to undergo pH-dependent interconversion between phenol-type and quinoid-type structures^32^. Therefore, as the assay was conducted at pH 7.4, both compounds are considered to predominantly exist in the quinoid form, and their metabolic behavior was nearly identical under these conditions.

#### Plasma protein binding

In plasma protein binding assays, the unbound fractions in human plasma were higher than those in mouse plasma, suggesting a greater proportion of free, pharmacologically active compound in human plasma. For the same reason as described above, the binding profiles of both compounds were also comparable under the assay conditions.

### 4. Berberrubine promotes mitochondrial biogenesis and increases mitochondrial protein expression

Next, we examined whether the increase in mitochondrial protein expression observed with quinoid-type berberrubine (Fig. 1g, h) was also evident with berberrubine acetic acid adduct. Similar to quinoid-type berberrubine, berberrubine acetic acid adduct significantly increased the protein level of MITOL (Fig. 3a). In addition, both compounds exhibited a dose-dependent trend in elevating the levels of other outer membrane proteins (Tom20, Mfn2), the matrix protein HSP60, and the inner membrane protein ATP5a. These effects were not observed with berberine. Furthermore, qRT-PCR analysis confirmed that both quinoid-type berberrubine and berberrubine acetic acid adduct increased the mRNA levels of MITOL and Tom20, exhibiting a dose-dependent trend (Fig. 3b). Consistent with the western blot results, these effects were not observed with berberine.

**Figure 3.**
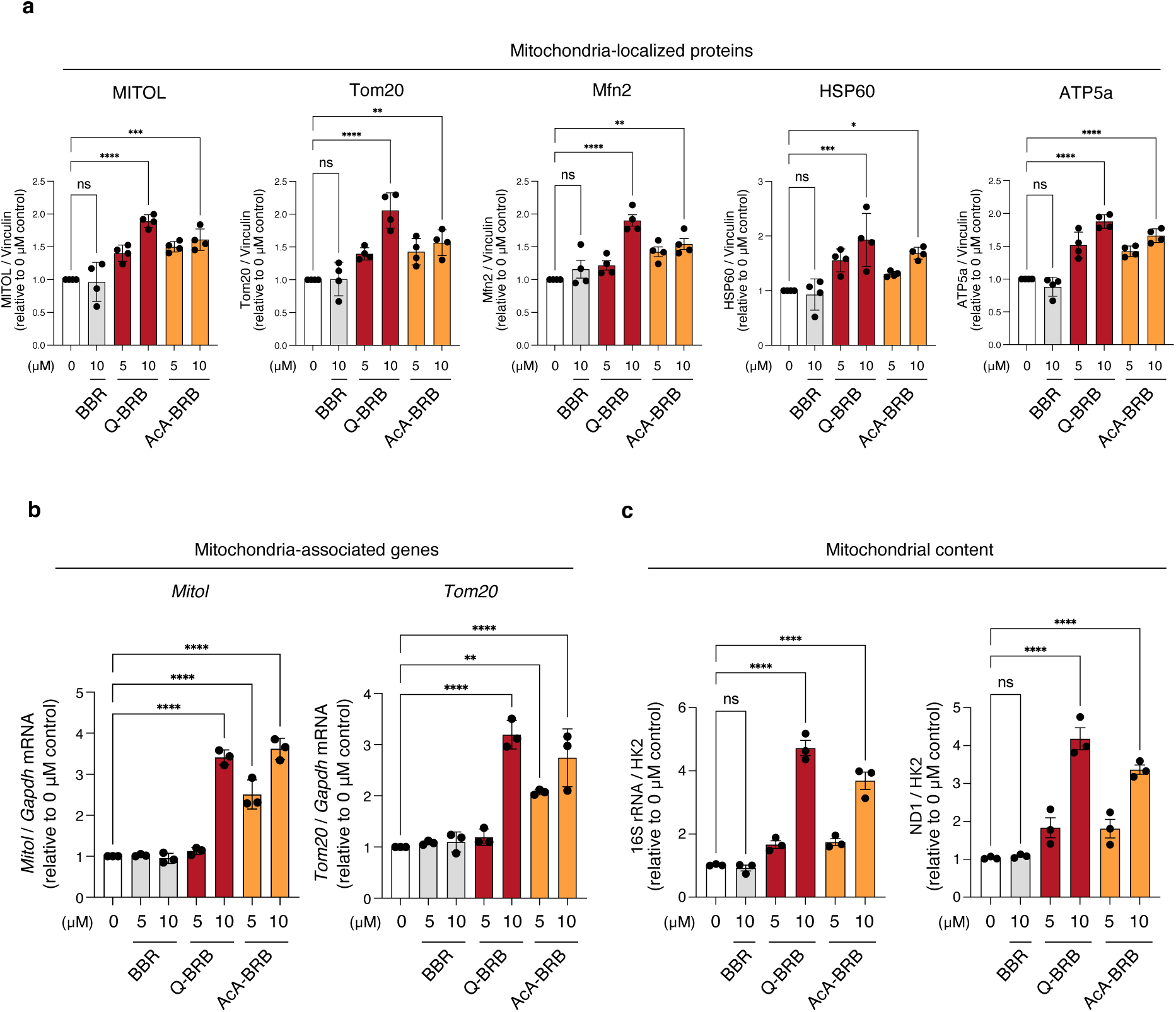
Enhancement of mitochondrial biogenesis and protein expression by berberrubine in C2C12 myoblasts. (a) C2C12 cells were treated with Q-BRB or AcA-BRB (5 μM, 10 μM), and the protein levels of mitochondrial-localized proteins (outer membrane: MITOL, Tom20, Mfn2; matrix: HSP60; inner membrane: ATP5a) were detected by Western blotting. Signal intensities were normalized to Vinculin and expressed as values relative to the 0 μM group (n = 4 per group). The 0 μM group indicates cells cultured without any additives, including solvents. (b) mRNA levels of MITOL and Tom20 were measured by qRT-PCR and normalized to GAPDH and expressed as fold change relative to the 0 μM group (n = 3 per group). (c) Mitochondrial content was quantified by real-time PCR based on the ratio of mtDNA (16S rRNA or ND1) to nuclear DNA (Hexokinase 2; HK2) and expressed as fold change relative to the 0 μM group (n = 3 per group). Statistical significance was determined using one-way ANOVA followed by Tukey’s HSD test. \**p* < 0.05; \*\**p* < 0.01; \*\*\**p* < 0.001; \*\*\*\**p* < 0.0001; ns, not significant.

Interestingly, both quinoid-type berberrubine and berberrubine acetic acid adduct significantly increased mitochondrial abundance at 10 μM (Fig. 3c), the concentration at which they also upregulated mitochondrial protein expression. Taken together, these findings indicate that berberrubine promotes mitochondrial biogenesis and enhances the expression of mitochondrial proteins at both mRNA and protein levels.

### 5. Berberrubine enhances mitochondrial elongation and mitochondrial respiration

Since MITOL regulates mitochondrial morphology through its E3 ubiquitin ligase activity, we examined mitochondrial morphological changes induced by berberrubine. As a result, mitochondrial elongation was significantly enhanced in C2C12 cells treated with berberrubine (Fig. 4a, b).

**Figure 4.**
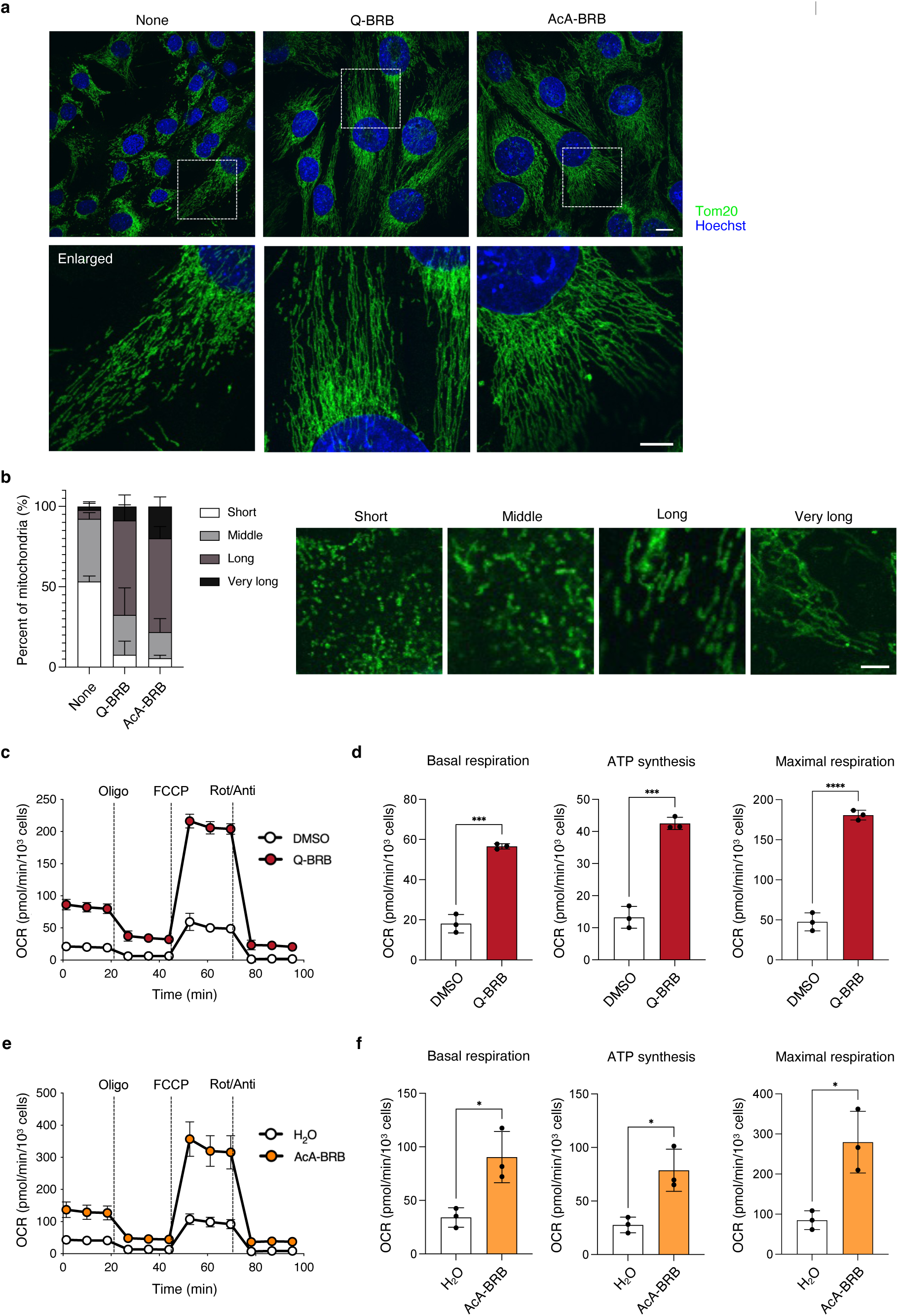
Mitochondrial elongation and increased oxygen consumption by berberrubine in C2C12 myoblasts. (a) C2C12 cells were treated with Q-BRB or AcA-BRB (10 μM) for 48 h, followed by immunostaining with an anti-Tom20 antibody to visualize mitochondrial morphology (green: anti-Tom20; blue: Hoechst). Representative images are shown in the upper panels. Enlarged views of the boxed regions are displayed in the lower panels. Scale bars: 20 μm (upper), 10 μm (lower). The None group indicates cells cultured without any additives, including solvents. (b) Mitochondrial length was quantified in 30 randomly selected cells per group using MiNA (Mitochondrial Network Analysis). Mitochondria were classified into four categories based on length: Short (≤1.855 μm), Middle (1.855–3.599 μm), Long (3.599–8.103 μm), and Very long (≥8.103 μm), and presented as the proportion of mitochondria in each category within each group. Representative images for each category are shown on the right. Scale bar: 10 μm. **(c, e)** C2C12 cells were treated with Q-BRB or AcA-BRB (30 μM) for 24 h, and mitochondrial oxygen consumption rate (OCR) was measured (n = 3 per group). **(d, f)** Basal respiration, ATP synthesis-linked respiration, and maximal respiration were calculated from the OCR data obtained in (c, e) after sequential administration of oligomycin (Oligo), FCCP, and rotenone/antimycin A (Rot/Anti). Statistical significance was determined by comparing each treatment group (Q-BRB or AcA-BRB) with the corresponding solvent control group (DMSO or H_2_O) using a two-sided unpaired Student’s *t*-test. \**p* < 0.05; \*\*\**p* < 0.001; \*\*\*\**p* < 0.0001.

To further assess mitochondrial function, we measured the mitochondrial oxygen consumption rate (OCR). In C2C12 cells treated with quinoid-type berberrubine or berberrubine acetic acid adduct, basal respiration, ATP synthesis-linked respiration, and maximal respiration were all significantly increased compared to the control group (Fig. 4c-f). These findings suggest that berberrubine enhances mitochondrial elongation and mitochondrial respiration.

### 6. Pharmacokinetic evaluation of quinoid-type berberrubine and berberrubine acetic acid adduct

Next, we evaluated the pharmacokinetics of quinoid-type berberrubine and berberrubine acetic acid adduct in mice (Suppl. Fig. 2a–c). Following oral administration at both low (10 mg/kg) and high (100 mg/kg) doses, both compounds reached their maximum plasma concentration (C_max_) within 0.25 hours, indicating rapid absorption in mice.

In both compounds, the C_max_ and area under the curve (AUC) values were over tenfold higher in the high-dose group compared to the low-dose group. Additionally, the half-life (T_1/2_) was prolonged and the total clearance (CL_tot_/F) was reduced in the high-dose group, suggesting saturation of the elimination process. The overall pharmacokinetic profiles were similar between the two compounds.

To further evaluate the pharmacokinetics profile under repeated oral administration conditions, mice were fed a 0.1% quinoid-type berberrubine mixed diet or given drinking water containing 0.125, 0.5, or 2.0 mg/mL berberrubine acetic acid adduct for 7 days. Additionally, a 28-day administration group was included for the 0.5 mg/mL dose. Plasma concentrations and tissue distribution were subsequently evaluated. The 2 mg/mL berberrubine acetic acid adduct group that received a 7-day administration exhibited significantly higher plasma berberrubine concentrations compared to all other groups (Suppl. Fig. 2d). Of note, the plasma concentrations observed in these repeated *in vivo* administrations were lower than the effective concentrations identified *in vitro*; nevertheless, as described in Fig. 5 and 6, pharmacological effects were clearly observed *in vivo*. These findings suggest that additional factors present *in vivo* may influence sensitivity to berberrubine, resulting in effective responses at lower concentrations than those observed *in vitro*.

**Figure 5.**
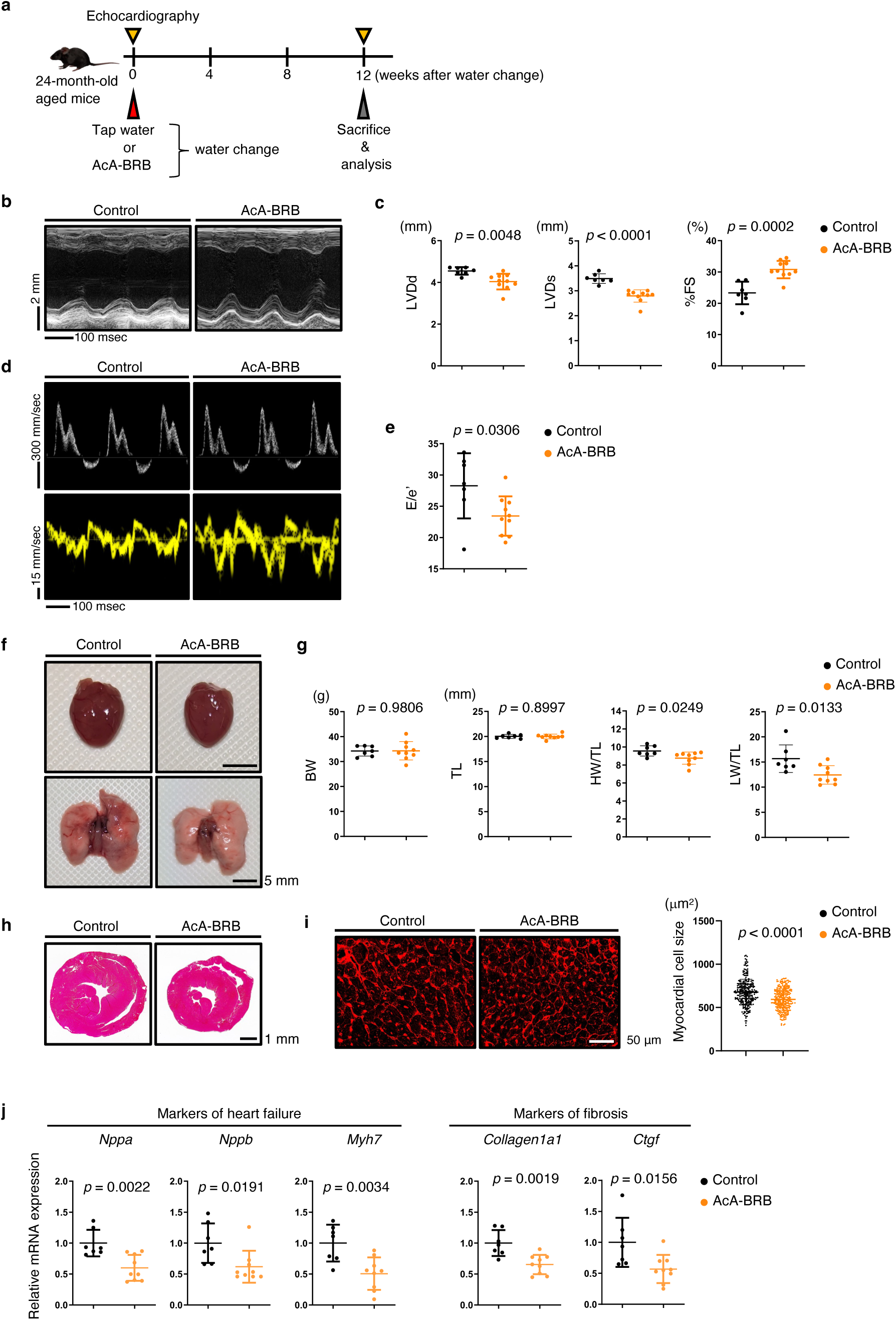
Cardioprotective effects of berberrubine in aged mice with age-related cardiac dysfunction. **(a)** Experimental scheme of berberrubine treatment in aged mice. **(b)** Representative M-mode echocardiography recordings in the indicated mice. **(c)** Echocardiographic measurements of left ventricular end-diastolic diameter (LVDd, mm), left ventricular end-systolic diameter (LVDs, mm), and percent fractional shortening (%FS) at 8 weeks after intervention (n = 7, control; n = 10, AcA-BRB). **(d)** Representative echocardiographic recordings of transmitral flow velocity, including early transmitral flow velocity (E), atrial systolic velocity (A) (top row), and early diastolic mitral annular velocity (e’) (bottom row) in the indicated groups. **(e)** E/e’ ratios in the indicated mice (n = 7, control; n = 10, AcA-BRB). **(f)** Gross appearance of whole hearts (top row; scale bar, 5 mm) and whole lungs (bottom row; scale bar, 5 mm) from the indicated mice. **(g)** Body weight (BW, g), tibial length (TL, mm), heart weight/tibial length ratio (HW/TL, mg/mm), and lung weight/tibial length ratio (LW/TL, mg/mm) in the indicated mice (n = 7, control; n = 9, AcA-BRB). **(h)** Hematoxylin-eosin (HE) staining of mid-portion heart sections from the indicated groups (scale bar, 1 mm). **(i)** Representative WGA-stained sections of the left ventricle indicating cardiomyocyte size (left; scale bar, 50 μm) and size distribution (right; n = 281 cells, control; n = 283 cells, AcA-BRB) of myocardial cells (μm^2^) in the indicated mice. **(j)** qRT-PCR analysis of heart failure markers (*Nppa*, *Nppb*, *Myh7*) and fibrosis markers (*Collagen1a1, Ctgf*) in the left ventricles of the indicated mice (n = 7, control; n = 9, AcA-BRB). Relative mRNA expression levels were normalized to 18S rRNA (*Rps18*) and presented as fold changes relative to the control group. Statistical significance was determined using a two-sided unpaired Student’s *t*-test (**c**, **e**, **g**, **i**, and **j**).

**Figure 6.**
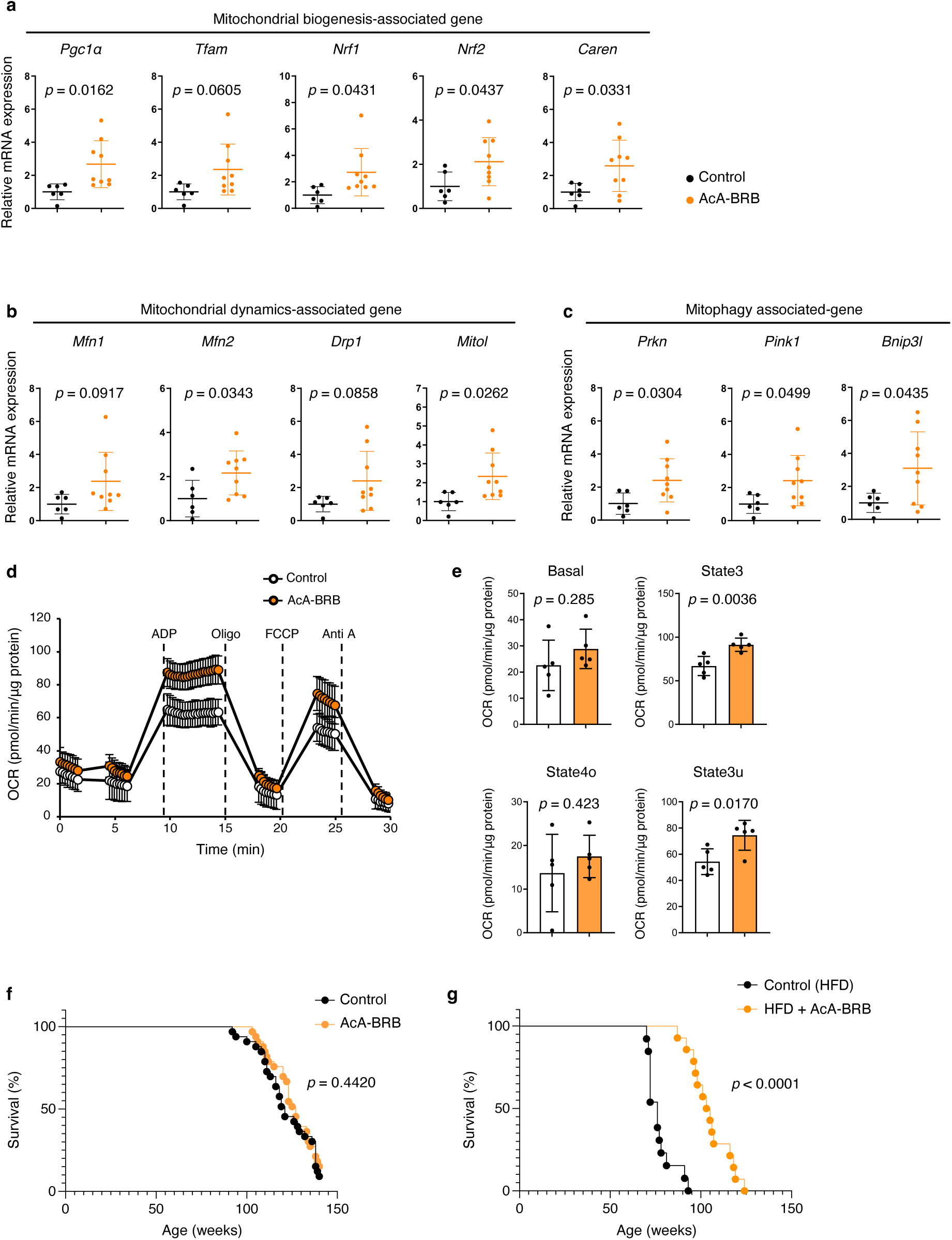
Effects of berberrubine on mitochondrial function and lifespan in aged mice with age-related cardiac dysfunction. (a-c) Relative expression of genes associated with mitochondrial (**a**) biogenesis, (**b**) dynamics, and (**c**) mitophagy in heart tissues of the indicated mice (n = 6, control; n = 9, AcA-BRB). (d) OCR analysis of mitochondria isolated from heart tissues of the indicated groups (n = 5 per group). (e) Quantification of basal respiration (Basal), ATP-linked respiration in the presence of ADP (State 3), resting respiration in the presence of oligomycin (State 4o), and maximal uncoupled respiration in the presence of FCCP (State 3u) (n = 5 per group) in the indicated groups. (f) Survival curve of the indicated groups (n = 33 per group). (g) Survival curve of the indicated groups (n = 13, control; n = 14, AcA-BRB). Statistical significance was determined using a two-sided unpaired Student’s *t*-test (**a**, **b**, **c**, **d**, and **e**) or the log-rank (Mantel-Cox) test (**f** and **g**).

Furthermore, tissue distribution analysis in the brain, heart, and skeletal muscle of this group revealed that berberrubine levels were exceptionally low in the brain, whereas a tendency toward higher concentrations was observed in the heart compared to other tissues (Suppl. Fig. 2e).

### 7. Berberrubine exerts protective effects against age-related cardiac dysfunction

We asked whether berberrubine administration in aged mice would ameliorate age-related cardiac dysfunction. To do so, we divided 24-month-old male C57BL/6N mice into two groups. One group received a standard diet with regular water, and the other received a standard diet with berberrubine-supplemented water. Based on preliminary experiments showing that aged mice refused to drink water with 2 mg/mL of berberrubine, we adjusted the concentration to 0.5 mg/mL to ensure free-drinking administration. After 12 weeks, we evaluated cardiac function (Fig. 5a). Echocardiographic analysis showed that age-related progression of cardiac left ventricular dilatation, systolic dysfunction and diastolic dysfunction was ameliorated in the berberrubine-treated group compared to control mice (Fig. 5 b-e). In addition, cardiac hypertrophy, as assessed by the HW/BW ratio and histological analysis, and cardiac function, as assessed by pulmonary congestion based on LW/BW ratio, were improved in berberrubine-treated mice relative to control mice (Fig. 5f-i). qRT-PCR analysis of heart tissue showed that upregulation of genes associated with heart failure and cardiac fibrosis was blocked in berberrubine-treated relative to control mice (Fig. 5j).

Interestingly, we confirmed that berberrubine treatment also significantly increased mitochondrial biogenesis-associated gene expression (Fig. 6a) as well as mitochondrial dynamics- and mitophagy-associated gene expression (Fig. 6b, c). OCR of mitochondria isolated from hearts of aged mice treated with berberrubine was improved compared to controls (Fig. 6d, e).

Blood pressure and heart rate were comparable in berberrubine-treated mice and control mice (Suppl. Fig. 3). Next, we asked whether long-term berberrubine treatment could prolong their lifespan. To address this, we evaluated the effect of long-term berberrubine administration on lifespan in 86-week-old aged mice compared with untreated groups. Notably, administration of 0.5 mg/mL berberrubine did not significantly affect the survival rate in this model (Fig. 6f). However, this finding suggests that berberrubine does not induce any adverse effects that could compromise survival in aged mice.

To further explore the impact of berberrubine on lifespan, we then examined its effect in a high-fat diet-induced aging model. Starting at 8 weeks of age, mice were fed a high-fat diet (HFD32). At 60 weeks old, the mice were divided into two groups: one group continued to receive tap water, while the other group was given berberrubine in drinking water (0.5 mg/mL). Lifespan was then compared between the two groups. As a result, the berberrubine-treated mice had a significantly prolonged lifespan compared with the control mice (Fig. 6g). These findings indicate that while berberrubine does not directly extend lifespan in aged mice with heart failure, it enhances cardiac function without increasing mortality risk and contributes to lifespan extension under metabolically stressful conditions and certain pathological states. Collectively, these results suggest that berberrubine improves age-related cardiac dysfunction and supports overall health, leading to a prolonged lifespan.

## Discussion

### 1. Potential of “Mitorubin” as a therapeutic agent for age-related cardiac dysfunction and its application to age-related diseases

In this study, we focused on mitochondrial ubiquitin ligase MITOL, a key enzyme that regulates mitochondrial morphology and function, and investigated the therapeutic effects of its pharmacological induction in age-related cardiac dysfunction. Our previous studies have demonstrated that MITOL expression declines with aging, and its depletion induces myocardial aging, leading to the onset of heart failure^18^. Furthermore, reduced MITOL expression has been implicated in neurodegenerative diseases such as Alzheimer’s and Parkinson’s disease^19,20^. Based on these findings, we screened compounds capable of increasing MITOL expression and identified that extracts from *Coptis japonica* and *Phellodendron amurense*, traditional herbal medicines, significantly increased MITOL mRNA and protein expression. Further analysis revealed that although berberine, the primary component of these extracts, did not directly activate mitochondria, its major metabolite, berberrubine, effectively induced MITOL expression. However, the poor water solubility of berberrubine posed a challenge. To address this, we synthesized a novel derivative, the berberrubine acetic acid adduct, which exhibits improved water solubility and allows for enhanced formulation flexibility and oral usability. We designated the group of mitochondria-activating berberrubine-based compounds, including this derivative, as “Mitorubin”.

To evaluate the physiological effects of Mitorubin, we administered it orally to 24-month-old aged mice. The results demonstrated that Mitorubin attenuated the progression of age-related cardiac dysfunction and improved cardiac function. Specifically, Mitorubin treatment suppressed left ventricular dilation and systolic dysfunction, as well as cardiac hypertrophy progression, indicative of reverse remodeling of the heart. Additionally, it increased the expression of mitochondria-associated genes and enhanced mitochondrial respiration. The pathology of age-related cardiac dysfunction involves mitochondrial dysfunction, disruption of mitochondrial dynamics, and increased oxidative stress. Our study suggests that Mitorubin suppresses the progression of age-related cardiac dysfunction by enhancing mitochondrial ATP production and normalizing mitochondrial dynamics.

Notably, Mitorubin-treated mice exhibited a significant extension of lifespan under metabolically stressful conditions, as observed in high-fat diet-fed mice. This finding suggests that Mitorubin may exert protective effects beyond cardiac function, potentially influencing systemic energy metabolism. Interestingly, a previous study has reported that berberrubine modulates glucose and lipid metabolism via gut microbiota^33^, indicating that its physiological effects may be mediated through multiple pathways *in vivo*.

In light of the observed enhancement of mitochondrial function in cardiac tissue and C2C12 myoblasts, it is plausible that Mitorubin may also exert beneficial effects in skeletal muscle. Since exercise tolerance is recognized as an important prognostic indicator for heart failure alongside conventional cardiac function indices in clinical practice^34^, we have initiated exploratory studies on the effects of Mitorubin on exercise performance in rodents and other species. Considering the well-documented association between heart failure and age-related conditions such as sarcopenia and frailty^35^, these studies are expected to provide insights into the potential therapeutic relevance of Mitorubin for age-associated muscular disorders.

The therapeutic potential of Mitorubin in neurodegenerative diseases remains another area warranting further investigation. Although Mitorubin exhibits lower brain permeability than other organs, its oral administration to mice led to an increase in MITOL expression in the brain (data not shown). Given that mitochondrial dysfunction plays a critical role in neurodegenerative diseases, we are currently conducting experiments to evaluate its efficacy in Alzheimer’s and Parkinson’s disease models.

### 2. Mechanism of Mitorubin action and its role in mitochondrial homeostasis

Elucidating the molecular mechanisms by which Mitorubin induces the expression of MITOL and other mitochondrial proteins remains a critical challenge, as these pathways have yet to be fully defined. Berberine has been reported to enhance mitochondrial biogenesis through the activation of the SIRT1-AMPK-PGC-1α pathway^36^ and to regulate mitochondrial quality control by promoting mitophagy^37^. Additionally, as berberine is known to accumulate in mitochondria^38^, Mitorubin may also specifically interact with mitochondrial proteins. Indeed, berberine is known to bind to the mitochondrial calcium uniporter (MCU) and inhibit calcium influx into mitochondria, thereby exerting cardioprotective effects against ischemia-reperfusion (I/R)-induced myocardial injury^39^. Given their structural similarity, an interaction between Mitorubin and MCU is conceivable. However, in our current study, berberine did not induce MITOL expression or increase mitochondrial content, suggesting that Mitorubin regulates mitochondrial homeostasis through a mechanism distinct from that of berberine.

Specifically, Mitorubin may interact with a mitochondrial protein other than MCU, and this unknown molecule could function as a sensor for mitochondrial depletion. This sensor molecule, upon activation or inactivation by Mitorubin, may transmit a signal to the nucleus via PGC-1α, thereby promoting mitochondrial biogenesis. In other words, Mitorubin may create a cellular state that mimics mitochondrial depletion via mitophagy, inducing a compensatory increase in mitochondrial proliferation. Supporting this hypothesis, an upregulation of mitophagy-related gene expression was observed in the hearts of Mitorubin-treated mice.

Recent studies have reported that the antidiabetic drug metformin induces mild mitochondrial stress by partially inhibiting respiratory complex I, which in turn promotes adaptive mitochondrial activation^40–42^. This phenomenon is considered a form of mitochondrial hormesis (mitohormesis). Hormesis describes a biological response in which exposure to low levels of stress enhances cellular resilience, leading to increased stress resistance^43^. In mitochondrial hormesis, mild mitochondrial stress activates antioxidant defenses and repair mechanisms, ultimately improving energy production^12,13^. Mitochondrial hormesis is also known to be induced by caloric restriction and physical exercise, both of which have been implicated in promoting longevity and extending healthspan^44–46^. However, the precise molecular mechanisms underlying mitochondrial hormesis remain incompletely understood, particularly how stress responses are relayed to transcription factors and signaling pathways that contribute to homeostasis. A recent review highlighted a mitohormetic response underlying the cytoprotective effects of berberine^47^. Elucidating the similarities and differences between mitohormetic mechanisms induced by berberine and Mitorubin represents an intriguing direction for future research. Emerging evidence suggests that higher-order assemblies of the mitochondrial electron transport chain, termed respiratory chain supercomplexes (respiratory supercomplexes), as key machinery for optimizing OXPHOS performance and potentially minimizing ROS production^48,49^. These supercomplexes facilitate more efficient electron flow between complexes and contribute to the structural stability of individual respiratory complexes. Age-associated changes in supercomplex organization have been reported and are thought to underlie mitochondrial dysfunction and age-related pathophysiological conditions^50^. Notably, our previous study demonstrated that cardiac-specific MITOL knockout mice exhibit impaired supercomplex formation, reduced myocardial ATP levels, and increased ROS production, indicating a critical role of MITOL in maintaining mitochondrial respiratory efficiency and redox balance^18^.

It remains to be clarified whether Mitorubin promotes the formation of respiratory supercomplexes through the upregulation of MITOL, and whether this process is mechanistically linked to the observed induction of mitochondrial biogenesis or occurs via independent pathways. Future studies will be essential to clarify how Mitorubin orchestrates both the quantity and functional capacity of mitochondria.

Furthermore, our preliminary observations suggest that the cellular effects of Mitorubin may vary depending on the metabolic state of the cell. In glycolysis-dependent cells, for instance, its effects appear to differ from those in OXPHOS-reliant cells, suggesting that metabolic dependence on OXPHOS may influence the cellular response to Mitorubin.

Given these insights, it is possible that Mitorubin engages a novel mitochondrial adaptation pathway distinct from that of metformin in activating MITOL. Further research is required to elucidate the detailed mechanisms involved. To address this, we are currently conducting target identification studies using drug affinity responsive target stability (DARTS) analysis and biotin-labeled probe assays to determine the mitochondrial binding partner of Mitorubin. Identification of this target protein would not only contribute to a deeper understanding of mitochondrial homeostasis but could also unveil novel therapeutic targets for drug development.

### 3. Future perspectives

The findings of this study suggest that Mitorubin holds promise as a potential therapeutic agent for age-related diseases. In both aged mice and those subjected to a high-fat diet, long-term administration of Mitorubin at 0.5 mg/mL did not lead to an increase in mortality; moreover, lifespan extension was observed in the latter group. These results suggest a favorable safety profile of Mitorubin under the present experimental conditions. Notably, recent studies have reported that high doses (up to100 mg/kg/day) of berberrubine can induce hepatic toxicity through metabolic activation and renal toxicity via distinct mechanisms in rodents^51–53^. These toxicities were observed following forced administration (intraperitoneal or oral gavage) of berberrubine chloride, in contrast to the *ad libitum* administration of a berberrubine acetic acid adduct used in our study, which differs in both compound properties and exposure profiles. Such differences may explain the absence of toxicity observed in our experimental setting. Nevertheless, it remains to be determined whether similar toxicity-related pathways could also be activated under Mitorubin treatment. Future studies should evaluate whether the effective dose and the toxic threshold are sufficiently separated to ensure safety, for example by determining the therapeutic index (TI), as this will be a key step toward clinical application.

To facilitate the clinical translation of Mitorubin, it is also necessary to clearly define its therapeutic indications. Age-related cardiac dysfunction, muscle atrophy, and neurodegenerative disorders are among the conditions in which mitochondrial dysfunction plays a significant role. Clarifying which pathological conditions respond to Mitorubin, and to what extent, will enable more precise therapeutic applications. Accordingly, further research using disease models is required to optimize dosing strategies, determine treatment durations, and design future clinical trials.

Mitochondrial dysfunction is a common pathological feature of many age-related diseases, and its preservation or restoration is believed to contribute to extended healthspan. Mitorubin has the potential to maintain mitochondrial homeostasis and optimize energy production, thereby mitigating disease progression. Based on the findings of this study, further research is needed to confirm its efficacy and safety for clinical application. Given the central role of mitochondrial dysfunction, Mitorubin may represent a novel therapeutic option. Continued studies will be essential to validate its potential and define appropriate safety parameters.

## Materials & Methods

### Chemical Synthesis

Quinoid-type berberrubine was synthesized by thermal decomposition of berberine chloride, which was purchased from Nacalai Tesque as “berberine hydrochloride” (product No. 04729-02; commercially referred to as such in the catalog, but structurally identical to berberine chloride based on the listed CAS number [141433-60-5]). Briefly, berberine chloride (100 g) was placed in a round-bottom flask and subjected to pyrolysis at 190 °C under reduced pressure (0.1 kPa) for 3 hours. The resulting reddish-black solid was allowed to cool to room temperature and collected as quinoid-type berberrubine (79.4 g) in 92% yield. To synthesize the berberrubine acetic acid adduct, quinoid-type berberrubine (70.0 g) was ground into a fine powder and suspended in glacial acetic acid (700 mL; Kanto Chemical). The mixture was stirred at room temperature (25 °C) for one hour. A yellow precipitate gradually formed during the reaction. The resulting suspension was filtered under vacuum using a Büchner funnel while washing with diethyl ether. The solid was then dried under reduced pressure at 40 °C for 8 hours to obtain a yellow powder (berberrubine acetic acid adduct) (107 g) in 88% yield. To prepare the berberrubine acetic acid salt, the yellow powder was further dried under reduced pressure at 60 °C for 6 hours. The resulting reddish-brown solid was collected and stored in a desiccator until use. All synthesized compounds were structurally characterized using ^1^H NMR and solid-state ^13^C NMR (CPMAS). The spectra confirmed the identities of each compound, particularly the amount of acetic acid molecules associated with berberrubine in the adduct and salt forms.

### NMR analysis

^1^H NMR spectra were recorded using a JEOL AM300 (^1^H, 300 MHz) spectrometer. Deuterated dimethyl sulfoxide (DMSO-*d*_6_, 2.49 ppm) was used as the internal standard for ^1^H NMR spectra. Solid-state ^13^C CPMAS NMR spectra were recorded using a JEOL ECA400 (^13^C, 100 MHz) spectrometer. Adamantane (29.472 ppm) was used as the external standard for ^13^C NMR spectra.

### Screening of MITOL-inducing compounds in cell-based assay

Human dermal fibroblasts (Kurabo) were seeded at 1.0 × 10^5^ cells/mL in FibroLife BM medium (Kurabo) supplemented with the provided supplements. A 0.5-mL aliquot of the cell suspension was dispensed into each well of a 12-well plate and incubated at 37°C under 5% CO_2_. After 24 hours, the medium was replaced with FibroLife BM medium (Kurabo) containing all provided supplements except fetal bovine serum (FBS) and fibroblast growth factor (FGF), and the cells were cultured for an additional 24 hours. The medium was then replaced with medium containing the test substances, and the cells were incubated for another 24 hours. Following incubation, the medium was removed, lysis buffer was added, and the cell lysate was collected.

Total RNA was extracted from the lysates using the RNeasy Mini Kit (Qiagen) according to the manufacturer’s instructions. cDNA was synthesized from the extracted RNA using PrimeScript RT Master Mix (Takara Bio). The mRNA expression levels of GAPDH and MITOL were measured using real-time PCR (SYBR Green method) with the StepOnePlus system (Thermo Fisher Scientific), and MITOL expression was normalized to GAPDH. The following primers (Takara Bio) were used: GAPDH (HA067812) and MITOL (HA192576). To assess the upregulating effects of the test substances on MITOL gene expression, the measured mRNA levels were compared with those of the control group (medium without test substances). The final concentrations of each test substance were as follows: rice bran fermented extract (4.0%), Phellodendron bark (*Phellodendron amurense*) extract (1.5%), Coptis root (*Coptis japonica*) extract (1.5%), β-estradiol (100 μM), 5-aminolevulinic acid (5-ALA) (10 mM), peach (*Prunus persica*) extract (2.11%), β-nicotinamide mononucleotide (β-NMN) (1 mM), riboflavin (200 μM), thiotaurine (5 mM), Tamogi-take mashroom (*Pleurotus cornucopiae*) extract (1%), MA-5 (1 mM), metformin (10 mM), urolithin A (500 μM), and niacinamide (5 mM).

### Mouse experiments for aging and herbal extract treatment

For age-related analysis of MITOL expression, male C57BL/6J mice aged 1, 10, and 18 months were euthanized, and heart tissues were collected. Tissues were homogenized in RIPA buffer supplemented with protease and phosphatase inhibitors, and total protein concentrations were determined using the BCA assay. Equal amounts of protein were subjected to SDS-PAGE and transferred to PVDF membranes for immunoblotting. A rabbit polyclonal anti-MITOL antibody was produced by immunizing rabbits with a synthetic peptide (GCKQQQYLRQAHRKILNYPEQEEA) corresponding to amino acids 257–279 of MITOL^14^.

Twelve-week-old male C57BL/6J mice were randomly divided into treatment groups (n = 5-6 per group). Herbal extracts and control solution (50% ethanol) were each diluted 1.5-fold with distilled water prior to administration. All solutions were administered intraperitoneally once daily for nine consecutive days at a dosage calculated to deliver ethanol at 2 g/kg body weight. One week after the final injection, mice were euthanized under isoflurane anesthesia and analyzed. Organs including the heart, liver, and kidney were rapidly harvested, washed in ice-cold PBS, snap-frozen in liquid nitrogen, and stored at −80°C until analysis. Protein extraction and immunoblotting were performed as described above.

The animal experiments described above were conducted by authors who were affiliated with Tokyo University of Pharmacy and Life Sciences at the time of the study, and the experimental protocols were approved by the university’s Animal Use Committee.

### Cell culture and compound treatment

Murine myoblast C2C12 cells were used for various cellular experiments in this study. Two different sources of C2C12 cells were used depending on the experiment: C2C12 cells obtained from RIKEN BRC (RCB0987) were used for gene expression, mtDNA quantification, and mitochondrial morphology analysis; C2C12 cells from ATCC (CRL-1772) were used for OCR analysis. Cells were cultured in Dulbecco’s Modified Eagle Medium (DMEM; Sigma-Aldrich or Nacalai Tesque), supplemented with 10% FBS and 1% penicillin–streptomycin. Cells were maintained in a humidified incubator at 37°C with 5% CO_2_.

For cellular experiments, berberine or quinoid-type berberrubine (Q-BRB) was dissolved in DMSO, and berberrubine acetic acid adduct (AcA-BRB) was dissolved in distilled H_2_O. Each compound was diluted in culture medium to the indicated final concentrations. The final concentration of DMSO did not exceed 0.1% (v/v) in any condition.

### Antibodies

The following antibodies were used: anti-MITOL (LS-C164034, LSBio); anti-α-Tubulin (T-9026, Sigma-Aldrich); anti-Mfn2 (sc-100560, Santa Cruz Biotechnology); anti-HSP60 (sc-136291, Santa Cruz Biotechnology); anti-ATP5a (ab14748, Abcam); anti-Tom20 (11802-1-AP, Proteintech); and anti-Vinculin (V9131, Sigma-Aldrich).

### Immunoblotting

Proteins in sample buffer were separated by SDS-PAGE and transferred to Immobilon-P PVDF membranes (IPVH00010, Millipore). Blots were probed with the indicated antibodies, and protein bands on the blot were visualized using Chemi-Lumi One L (07880-70, Nacalai Tesque) or Immobilon Western Chemiluminescent HRP Substrate (WBKLS0500, Millipore). Band images were captured using a LuminoGraph I imager (WSE-6100, ATTO). Relative band intensities were quantified using Fiji/ImageJ software^54^.

### qRT-PCR analysis of C2C12 cells

Total RNA was extracted from C2C12 cells using an RNA extraction kit (RNeasy Mini Kit, Qiagen). After washing with PBS, cells were collected by trypsinization and centrifugation at 500 × *g* for 5 minutes, and the pellet was lysed according to the manufacturer’s instructions. The extracted RNA was reverse-transcribed into cDNA using a reverse transcription kit (ReverTra Ace, TOYOBO). Quantitative real-time PCR was performed using a qPCR master mix (KAPA SYBR FAST qPCR Master Mix (2X) Kit, KAPABIOSYSTEMS) and a thermal cycler (StepOnePlus, Applied Biosystems). Gene expression levels were normalized to GAPDH. Primer sets used for qRT-PCR are listed in Supplementary Table 1.

### Quantification of mitochondrial DNA

To assess mitochondrial DNA (mtDNA) content, the mtDNA/nuclear DNA (nDNA) ratio was determined by extracting DNA from C2C12 cells using the NucleoSpin Tissue Kit (Takara Bio), followed by real-time quantitative PCR analysis with specific primer sets for mtDNA (16S rRNA or ND1) and nDNA (Hexokinase 2; HK2) (Supplementary Table 2). Real-time quantitative PCR was performed using TB Green Premix Ex Taq II (Takara Bio) and a Thermal Cycler Dice Real Time System (Takara Bio). The analysis of mtDNA/nDNA ratios was conducted using the ΔΔCt method as previously described^55^.

### Mitochondrial morphology analysis

Cells were fixed with 4% paraformaldehyde in PBS for 15 minutes at 37 °C, washed three times with PBS, and permeabilized with 0.1% Triton X-100 in PBS for 10 minutes at room temperature (RT). After three additional washes with PBS, cells were blocked with 10% bovine serum albumin (BSA) in PBS for 10 minutes at RT. Cells were incubated with anti-Tom20 primary antibody (1:750 dilution, 11802-1-AP, Proteintech) in PBS containing 5% BSA for 1 hour at RT, followed by three PBS washes. Hoechst 33342 (1:2,000 dilution) and Alexa Fluor-conjugated secondary antibody (1:500 dilution, A11029, Invitrogen) in PBS containing 5% BSA were applied for 30 minutes at RT.

After three final PBS washes, cells were mounted with Fluorescent Mounting Medium (S3023, Dako), and imaged using an FV3000 confocal laser scanning microscope (Olympus). Images were acquired as Z-stacks with 0.2 μm intervals and reconstructed into maximum intensity projections using Fiji/ImageJ software.

Mitochondrial morphology was quantified using the MiNA plugin for Fiji (https://imagej.net/plugins/mina). Thirty cells per group were randomly selected for analysis. Average mitochondrial length and other morphological parameters were calculated according to the plugin’s standard analysis pipeline.

### Oxygen consumption analysis of C2C12 cells

The oxygen consumption rate (OCR) of C2C12 myoblasts was measured using a Seahorse XFe24 Analyzer (Agilent Technologies) and the Seahorse XF Cell Mito Stress Test Kit (Agilent Technologies), according to the manufacturer’s protocol. Cells were seeded at a density of 1.2 × 10^4^ cells per well in an Agilent Seahorse XF24 Cell Culture Microplate (Agilent Technologies). After 24 hours, cells were treated with berberrubine or the corresponding vehicle (DMSO or H_2_O) as indicated in the figure legends for 24 hours. One hour prior to the assay, the culture medium was replaced with Seahorse XF Base Medium supplemented with 1 mM sodium pyruvate, 2 mM glutamine, and 10 mM glucose (pH 7.4), and cells were equilibrated in a non-CO_2_ incubator at 37°C. The assay was performed at 37°C, and OCR was measured at baseline and after sequential injections of the following compounds: oligomycin (1 μM), FCCP (0.5 μM), and a mixture of rotenone and antimycin A (0.5 μM each). Basal respiration, ATP synthesis-linked respiration, and maximal respiration were calculated as follows: basal respiration = baseline OCR − non-mitochondrial respiration; ATP-linked respiration = baseline OCR − OCR after oligomycin injection; maximal respiration = OCR after FCCP injection − non-mitochondrial respiration.

### In vitro ADME assay

#### PAMPA assay

To determine the passive membrane diffusion rates, Corning Gentest Pre-coated PAMPA Plate System was used. The acceptor plate was prepared by adding 200 μL of 0.1 M phosphate buffer (pH 4.5-8.6) supplemented with 5% DMSO to each well, and then 300 μL of 30 μM compounds in 0.1 M phosphate buffer (pH 4.5-8.6) with 5% DMSO were added to the donor wells. The acceptor plate was then placed on top of the donor plate and incubated at 37 °C for 4 hours without agitation. After the incubation, the plates were separated and the solutions from each well of both the acceptor plate and the donor plate were transferred to 96-well plates and mixed with acetonitrile. The final concentrations of compounds in both donor and acceptor wells, as well as the concentrations of the initial donor solutions, were analyzed using liquid chromatography-tandem mass spectrometry (LC-MS/MS). The permeability of the compounds was calculated as described in a previous study^56^. Antipyrine (100 μM; 100% gastrointestinal absorption in humans), metoprolol (500 μM; 95%) and sulfasalazine (500 μM; 13%) were used as reference compounds. The permeabilities of antipyrine, metoprolol and sulfasalazine were 24, 3.0 and 0.084 × 10^-6^ cm/s, respectively. Two technical replicates were performed.

#### Hepatic microsomal stability assay

Disappearance of the parent compound over time was measured by using the amount of drug at time zero as a reference. After 5 minutes of preincubation, 1 mM NADPH (final concentration; the same applies to the following) was added to a mixture containing 1 μM of the compound, 0.2 mg/mL of human or mouse liver microsomes (Sekisui XenoTech), 1 mM EDTA and 0.1 M phosphate buffer (pH 7.4). The mixture solution was incubated at 37 °C for 30 minutes with rotation at 60 rpm. An aliquot of 50 μL of the incubation mixture was added to 250 μL of chilled acetonitrile/internal standard (IS, methyltestosterone). After centrifugation at 3,150 × *g* for 15 minutes at 4 °C, the supernatants were analyzed by LC-MS/MS. Hepatic microsomal stability (mL/min/kg, CL_int_) was calculated according to a previous report^57^, using 48.8 (human) or 45.4 (mouse) mg MS protein/g liver and 25.7 (human) or 87.5 (mouse) g liver/kg body weight as scaling factors. Two technical replicates were performed.

#### Determination of the unbound fraction in human or mouse plasma

An equilibrium dialysis apparatus was used to determine the unbound fraction for each compound in human or mouse plasma. High Throughput Dialysis Model HTD96b and Dialysis Membrane Strips MWCO 12-14 kDa (HTDialysis) were used. Plasma was spiked with the test compound (1 μM), and 150 μL aliquots were loaded into the apparatus and dialyzed versus 150 μL of 0.1 M phosphate buffer (pH 7.4) at 37 °C for 6 hours by rotation at 80 rpm. The unbound fraction was calculated as the ratio of receiver side (buffer) to donor side (plasma) concentrations. Two technical replicates were performed.

### In vivo pharmacokinetics assay

All mouse experiments were approved by and conducted in accordance with the guidelines of the Animal Experiment Committee of the University of Tokyo (permission number: P29-24). The compounds were suspended in 0.5% methylcellulose and orally administered to 8-week-old male C57BL/6N mice (Oriental Yeast) following an overnight fast. The dosage of the compound was 10 or 100 mg/10 mL/kg. Serial blood samples (<50 μL per time point) were collected from the caudal vein of each mouse at 15 and 30 minutes, and at 1, 2, 4, 6 and 24 hours post-dose using EM MYSTAR Hematocrit Capillary treated with heparin (AS ONE). Plasma was obtained by centrifugation of the blood samples. After the final blood sampling at 24 hours, the mice were humanely euthanized.

In the repeated administration study, animals were treated according to the group allocations as described in the main text and corresponding to Supplementary Figure 2d. Compounds were administered orally either via drinking water or mixed feed for 7 or 28 days, and plasma, brain, heart, and skeletal muscle samples were collected at the final time point (8:00-10:00 a.m.) on the day of sacrifice (n = 4 per group). Blood was drawn from the abdominal vena cava under terminal anesthesia and plasma was separated by centrifugation using heparinized tubes. Tissue samples were homogenized with four volumes of phosphate-buffered saline. Plasma and tissue homogenate samples were precipitated with acetonitrile containing an internal standard (IS) at a volume fourfold or greater than that of the sample, and then centrifuged at 15,000 × *g* for 10 minutes at 4 °C. The resulting supernatants were analyzed by LC-MS/MS.

Before pharmacokinetic analyses, plasma concentrations were averaged at each time point. Concentrations below the limit of quantitation were excluded from the analysis. Pharmacokinetic analyses were conducted using PhoenixWinNonlin version 8.3 (Pharsight Corp). The maximum plasma concentrations (C_max_) and times to achieve maximum plasma concentrations (T_max_) were obtained directly from the plasma concentration-time curves. The area under the concentration-time curve from zero to infinity (AUC_∞_) was obtained using the linear trapezoidal method in the noncompartmental analysis module. The software was allowed to determine the best-fit line for the elimination slope for all calculations. Other parameters such as elimination half-life period (t_1/2_), the apparent total body clearance CL_tot_/F, and volume of distribution V_dz_/F were further calculated.

### LC-MS/MS

An LCMS-8060 instrument equipped with a Shimadzu Nexera series LC system (Shimadzu) was used. Berberrubine and IS were analyzed in multiple reaction monitoring mode under electronspray ionization conditions. The analytical column used was a CAPCELL PAK C18 MGIII (3 μm × 2.0 mm ID × 35 mm; OSAKA SODA) at 50 °C. The gradient mobile phase consisted of 0.1% formic acid in water (mobile phase A) and 0.1% formic acid in acetonitrile (mobile phase B) at a total flow rate of 1 mL/min. The initial mobile phase composition was 10% B, which was held constant for 0.5 minute, increased in a linear fashion to 90% B over 1 minute, then held constant for 0.8 minute, and finally brought back to the initial condition of 10% B over 0.01 minute and reequilibrated for 1 minute. The transitions (precursor ion > product ion) of berberrubine and IS (methyl testosterone) are 322.11 > 307.1 and 303.1 > 109.1 (positive), respectively.

### Experimental animals for cardiac and survival analysis

All experimental procedures were approved by the Ethics Review Committee for Animal Experimentation at Kumamoto University, Kumamoto, Japan (approval No. A 2024-083). Procedures followed the guidelines and recommendations outlined by the ILAR Guide for the Care and Use of Laboratory Animals (8th Edition, 2011). Prior to tissue collection, mice were euthanized under isoflurane anesthesia, followed by cervical dislocation to ensure rapid euthanasia, a procedure also employed during sacrifice for tissue harvesting. Euthanasia was performed when animals met the established criteria for humane endpoints, specifically a body weight loss exceeding 20% over a 3-day period or 25% over a 1-week period, in accordance with the Kumamoto University Ethics Review Committee for Animal Experimentation above. All animals fed a normal diet (CE-2, CLEA) were randomly assigned to two groups: tap water only and tap water supplemented with AcA-BRB (0.5 mg/mL). Mice were housed in an animal facility with lighting cycles automatically controlled (12 h light/12 h dark), maintained at a stable temperature of 22 ± 2 °C and a relative humidity of 40–80%. C57BL/6NJcl wild-type (WT) male mice aged 2, 20, and 24 months were used in this study.

### High fat diet feeding in mice

For survival analysis under metabolic stress conditions, two-month-old C57BL/6NJcl male mice fed a high-fat diet (HFD 32: water 6.2%, protein 25.5%, fat 32.0%, ash 4.0%, carbohydrate 29.4%, fiber 2.9%, providing 507.6 kcal/100 g; CLEA) were randomly assigned to two groups: tap water only and tap water supplemented with AcA-BRB (0.5 mg/mL).

### qRT-PCR analysis of cardiac tissue

Total RNA was extracted using a RNeasy Mini Kit (Qiagen). DNase-treated RNA was reverse-transcribed using a PrimeScript RT reagent Kit (Takara Bio). Heart tissue was homogenized using a multi-beads shocker (Yasui Kikai). Real-time quantitative RT-PCR was performed using TB Green Premix Ex Taq II (Takara Bio), and a Thermal Cycler Dice Real Time System (Takara Bio). Relative transcript abundance was normalized to 18S rRNA levels. Primer sets used for RT-PCR are listed in Supplementary Table 3.

### Echocardiographic Analysis

Mice were preconditioned by chest hair removal with a topical depilatory (FUJIFILM VisualSonics), anesthetized with 1.5–2.5% isoflurane administered via inhalation, and maintained in a supine position on a dedicated animal handling platform with limbs immobilized for electrocardiogram gating during imaging. Body temperature was kept constant by relaying signals from a rectal probe to a heating pad, while heart and respiratory rates were continuously monitored. Transthoracic echocardiography was performed using a high-frequency ultrasound system dedicated to small-animal imaging (VisualSonics Vevo 3100, FUJIFILM VisualSonics) using a MS 400 linear array transducer (18–38 MHz). M-mode recording was performed at the midventricular level. All images were analyzed using dedicated software (Vevo LAB version 5.7.1). LV wall thickness and internal cavity diameters at diastole (LVID;d) and systole (LVID;s) were measured. Percent LV fractional shortening (%FS) was calculated from M-mode measurements. To analyze cardiac diastolic function, trans-mitral flow velocity patterns and early diastolic mitral annular velocity (e’) were measured in the four-chamber view to calculate the ratio of early trans-mitral flow velocity (E) to e’ (E/e’). All procedures were performed under double-blind conditions with regard to genotype or treatment.

### Isolation of mouse heart mitochondria

Mouse heart tissue was homogenized in mitochondrial isolation buffer (210 mM mannitol, 70 mM sucrose, 5 mM HEPES-KOH, 1 mM EGTA, 0.5% fatty acid-free BSA, pH 7.2) using a glass-teflon homogenizer. Homogenates were centrifuged at 27,000 × *g* for 10 minutes at 4 °C, and the resulting pellets were resuspended in mitochondrial isolation buffer. Samples were centrifuged at 500 × *g* for 5 minutes at 4 °C, and the supernatants were collected and centrifuged at 10,000 × *g* for 5 minutes at 4 °C. The pellet was then resuspended in mitochondrial isolation buffer and centrifuged for 5 minutes at 10,000 × *g* at 4 °C. The resulting pellets were suspended in BSA-free mitochondrial isolation buffer, and mitochondrial protein concentration was determined using the Bio-Rad Protein Assay Dye Reagent (Bio-Rad Laboratories). Samples were then incubated with an equal volume of mitochondrial isolation buffer and subjected to oxygen consumption analysis.

### Oxygen consumption analysis of isolated cardiac mitochondria

Isolated mitochondria were diluted with mitochondrial assay solution (220 mM mannitol, 70 mM sucrose, 2 mM HEPES-KOH, 1 mM EGTA, 10 mM KH_2_PO_4_, 5 mM MgCl_2_, 0.2% fatty acid-free BSA, 2 mM malate, 10 mM pyruvate, pH 7.2) to 80 μg/mL. The diluted mitochondria were loaded at 50 μL per well into 24-well Seahorse assay plates (Seahorse Bioscience) and centrifuged at 2,000 × *g* for 20 minutes at 4 °C. After centrifugation, 450 μL mitochondrial assay solution was added to each well, and plates were incubated for 8 minutes at 37 °C. To assess mitochondrial respiratory function, the following compounds (final concentrations) were sequentially injected into each well: ADP (4 mM), oligomycin (2 μM), FCCP (4 μM), and antimycin A (4 μM). OCR was measured under basal conditions and after each injection using an XFe24 extracellular flux analyzer (Seahorse Bioscience).

### Statistical analysis

Data are presented as mean ± standard error of the mean (SEM), unless otherwise indicated. Statistical analyses were performed using GraphPad Prism 9 or GraphPad Prism 10 software (GraphPad Software). The specific statistical methods used for each dataset are described in the corresponding figure legends. P values are either directly shown above the bars or indicated using the following notation: \**p* < 0.05; \*\**p* < 0.01; \*\*\**p* < 0.001; \*\*\*\**p* < 0.0001; ns, not significant.

## Acknowledgments

We thank Taisho Pharmaceutical Co., Ltd. (Tokyo, Japan) for their significant contributions to the selection of candidate compounds for MITOL induction, the provision of information regarding the panel of herbal extracts and bioactive compounds, and support in related experimental procedures. We are also grateful to Ms. Eri Nakashima, Mr. Yasuyuki Kure, and Ms. Noriko Nakaya for their invaluable technical assistance with the animal experiments and LC-MS/MS analysis. This work was supported in part by the Japan Agency for Medical Research and Development (AMED) [Grant Numbers JP24ama121051 (H. Kusuhara) and JP24ama121053 (K. Kanamitsu) under the Research Support Project for Life Science and Drug Discovery (BINDS); JP24gm1710006h0002 (Y. Oike) under the Core Research for Evolutional Science and Technology (CREST) Program; and DNW-24003 (S. Yanagi) under the Drug Discovery Booster Program]. This study was also supported by the Ministry of Education, Culture, Sports, Science and Technology (MEXT) and the Japan Society for the Promotion of Science (JSPS) KAKENHI [Grant Numbers 23K07511 (M. Sato), 22K11727 (D. Torigoe), 24K21297 (S. Inoue), 21H04825 (Y. Oike), and 23H02691 and 23K27382 (S. Yanagi)]. Additional support was provided by the Takeda Science Foundation (2024) [M. Sato, T. Kadomatsu, and Y. Oike] and the Japan Geriatrics Society (JGS) Research Grant Award in Geriatrics and Gerontology (2024) [M. Sato].

## Authors’ contributions

M. Sato, S. Inoue, H. Abe, Y. Oike, and S. Yanagi conceived the project and designed the study. T. Tokuyama contributed to chemical screening and compound selection. A. Yokosuka, Y. Mimaki, and H. Abe conducted chemical synthesis and analysis. M. Sato, D. Torigoe, T. Kadomatsu, E. Kanai, T. Hamano, and H. Hirata performed *in vivo* experiments and contributed to the interpretation and discussion of the results. D. Tanabu, K. Taniwaka, Y. Suzuki, and T. Takeiwa carried out molecular and cellular experiments. K. Kanamitsu and H. Kusuhara performed pharmacokinetic and ADME analyses. Data analysis and interpretation were conducted by M. Sato, D. Torigoe, D. Tanabu, Y. Ogata, and I. Shiiba. M. Sato, D. Torigoe, D. Tanabu, Y. Ogata, R. Inatome, and S. Yanagi contributed to manuscript preparation and editing. H. Abe, Y. Oike, and S. Yanagi supervised the study. Funding was acquired by M. Sato, D. Torigoe, S. Inoue, K. Kanamitsu, H. Kusuhara, Y. Oike, and S. Yanagi.

---

**Supplementary Table 1.**
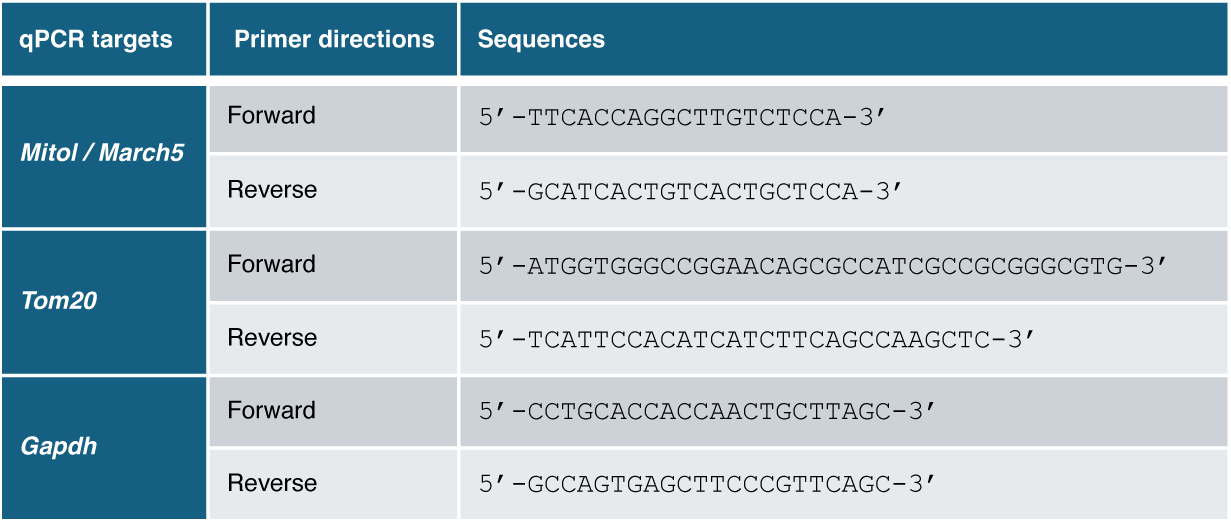
Primer pairs used for quantitative RT-PCR analysis in C2C12 cells.

**Supplementary Table 2.**
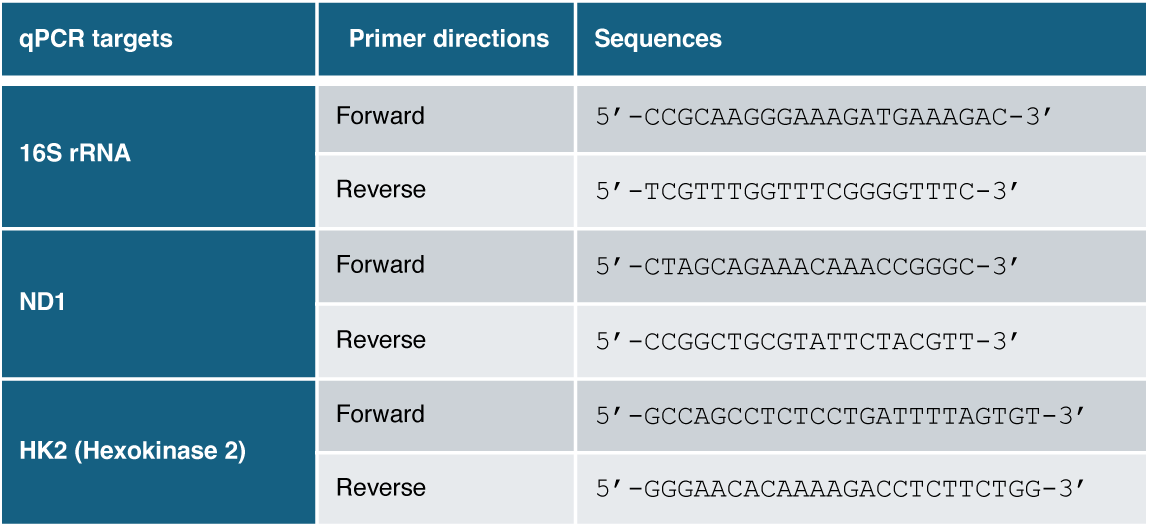
Primer pairs used to quantify mitochondrial DNA content.

**Supplementary Table 3.**
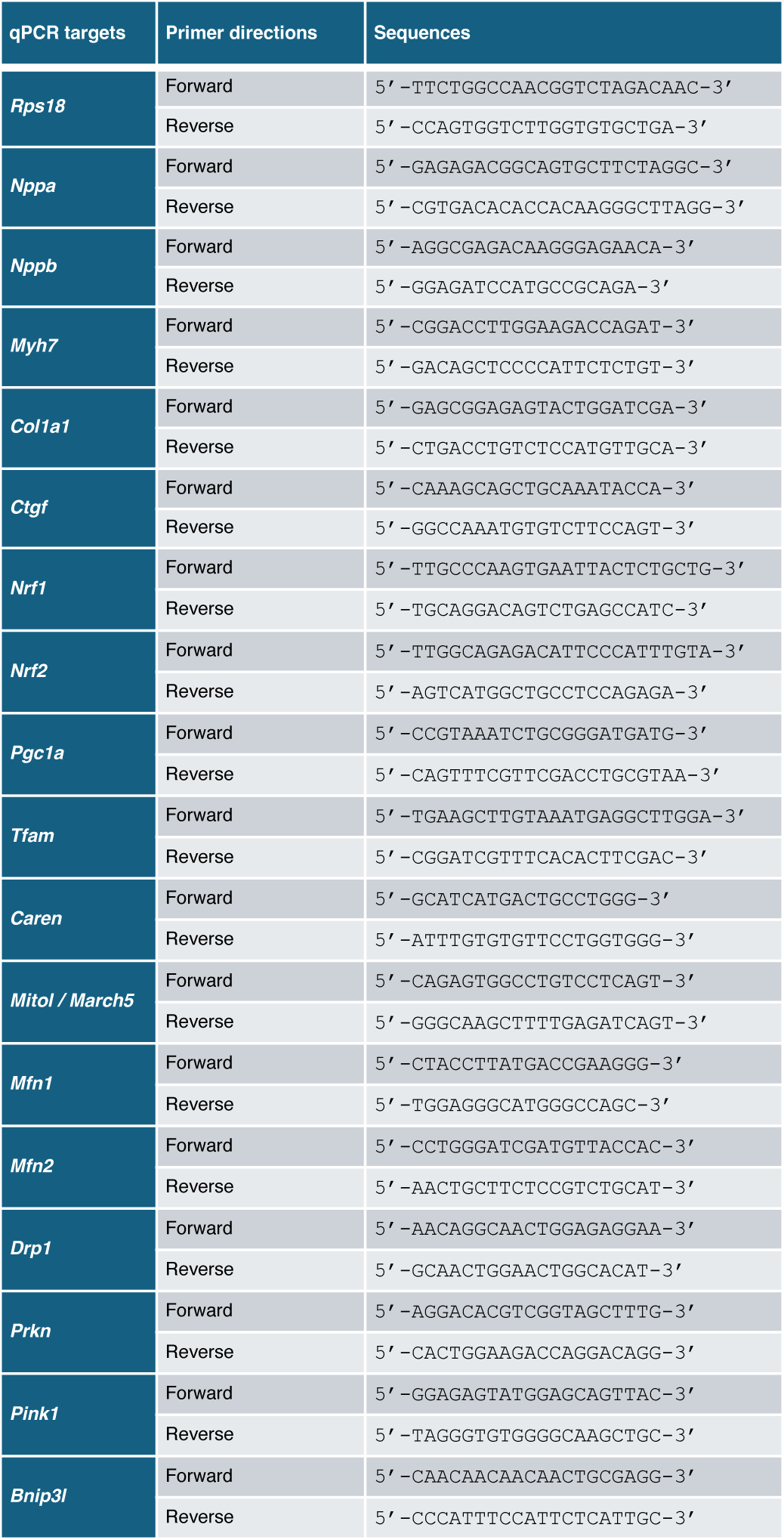
Primer pairs used for qRT-PCR analysis in mouse heart samples.

**Supplementary Figure 1.**
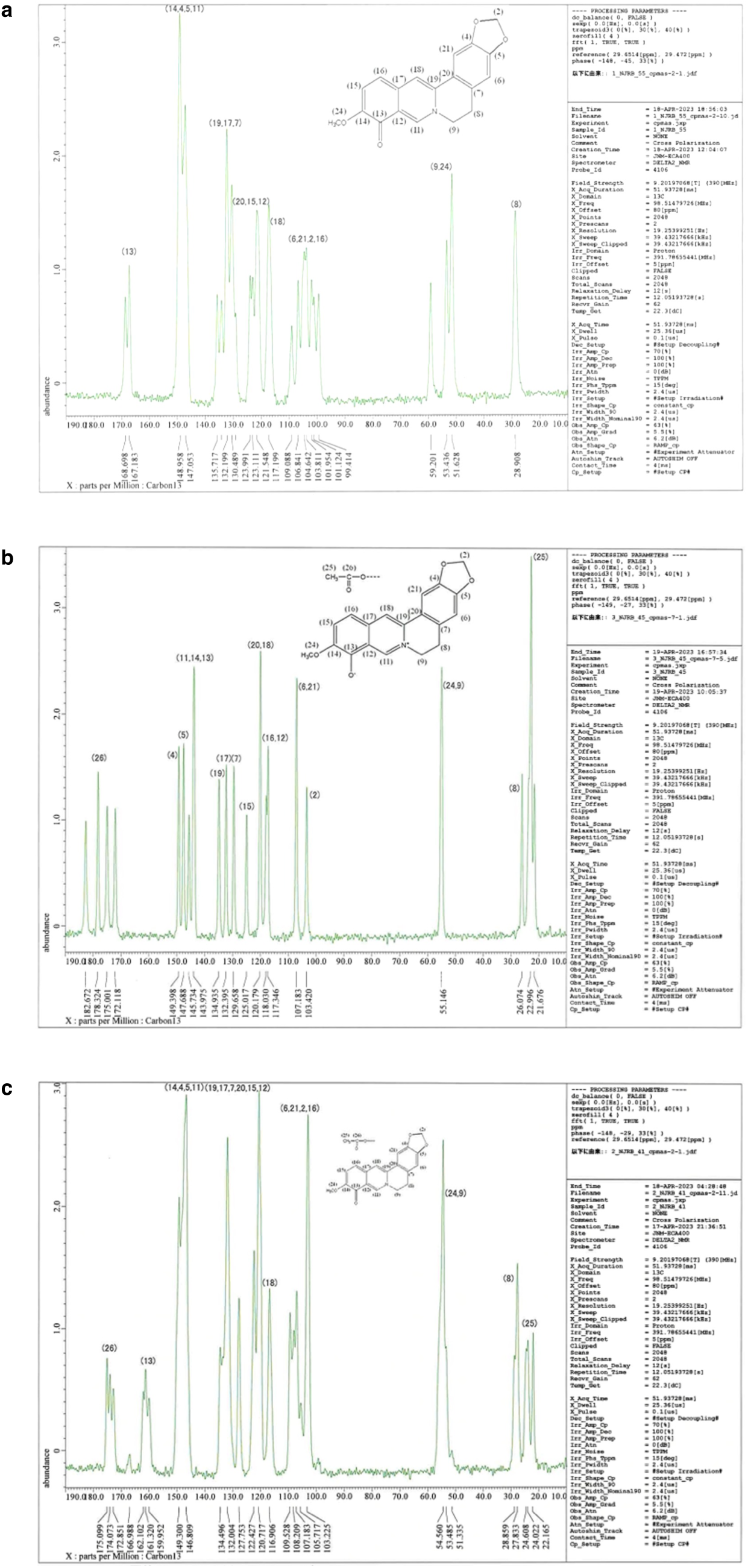
Structural analysis of quinoid-type berberrubine, berberrubine acetic acid adduct, and berberrubine acetic acid salt. Solid-state ^13^C-NMR spectra of the berberine-derived compounds were obtained using the CPMAS method. The spectra are presented from top to bottom: **(a)** reddish-black solid (quinoid-type berberrubine), **(b)** reddish-brown solid (berberrubine acetic acid salt), and **(c)** yellow solid (berberrubine acetic acid adduct).

**Supplementary Figure 2.**
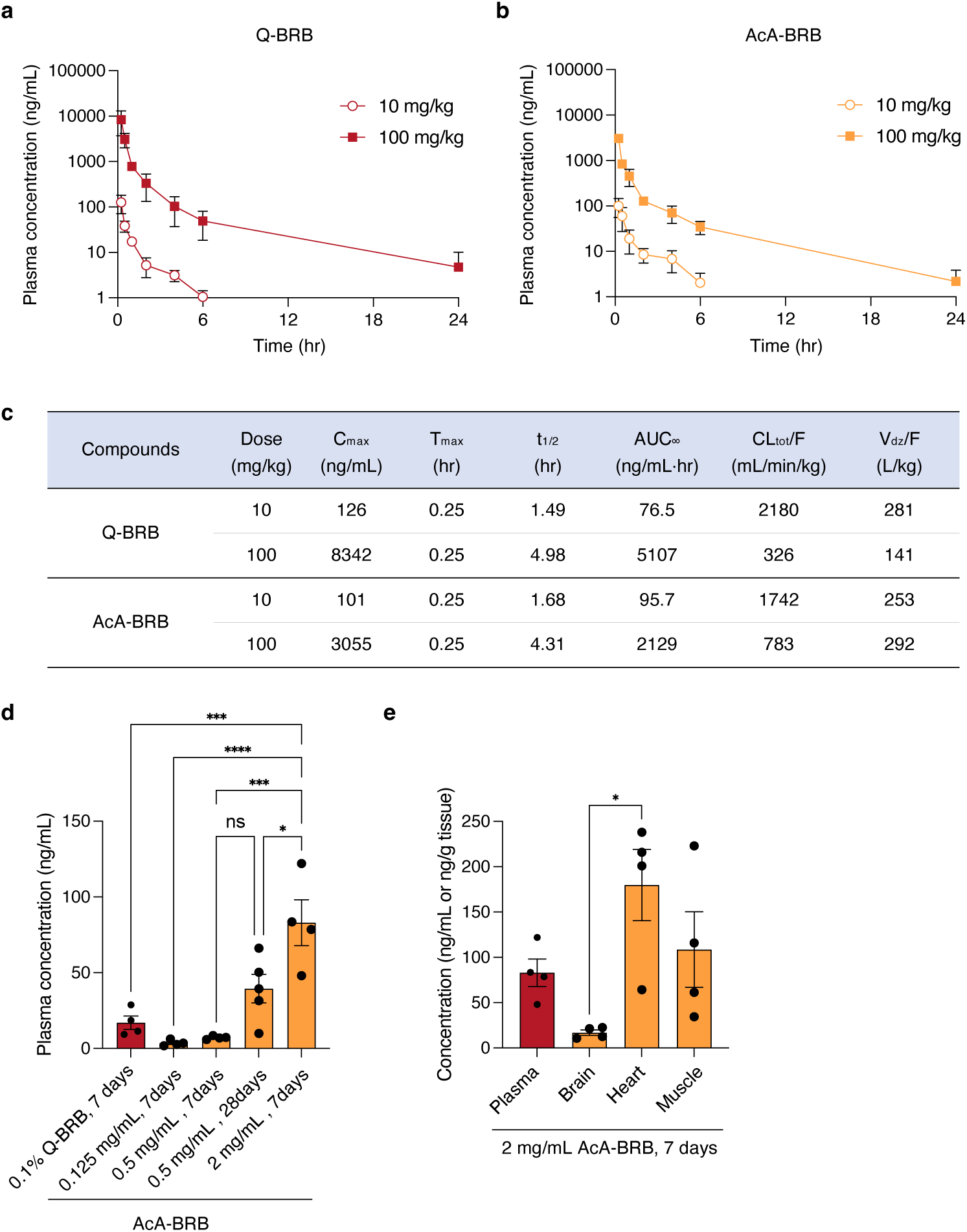
Pharmacokinetic evaluation of quinoid-type berberrubine and berberrubine acetic acid adduct in mice. (a, **b)** Plasma concentration profiles of quinoid-type berberrubine (Q-BRB, **a**) and berberrubine acetic acid adduct (AcA-BRB, **b**) following a single oral gavage administration at low (10 mg/kg) and high (100 mg/kg) doses (n = 3 per group). Data are presented as mean ± standard deviation (SD). (c) Pharmacokinetic parameters, including maximum plasma concentration (C_max_), time to reach Cmax (T_max_), elimination half-life (t_1/2_), area under the plasma concentration-time curve (AUC_∞_), apparent total clearance (CL_tot_/F), and volume of distribution (V_dz_/F), for Q-BRB and AcA-BRB at each dose. (d) Plasma berberrubine concentrations in mice after 7-day administration of either 0.1% Q-BRB mixed in diet or AcA-BRB in drinking water at concentrations of 0.125, 0.5, and 2.0 mg/mL (n = 4 per group), including an additional 28-day group for the 0.5 mg/mL dose (n = 5). (e) Plasma and tissue distribution of berberrubine in the brain, heart, and skeletal muscle of mice after 7-day administration of 2 mg/mL AcA-BRB in drinking water (n = 4 per group). Statistical significance was determined using one-way ANOVA followed by Tukey’s HSD test (**d**) and a two-sided unpaired Student’s *t*-test (**e**). \**p* < 0.05; \*\**p* < 0.01; \*\*\**p* < 0.001; ns, not significant.

**Supplementary Figure 3.**
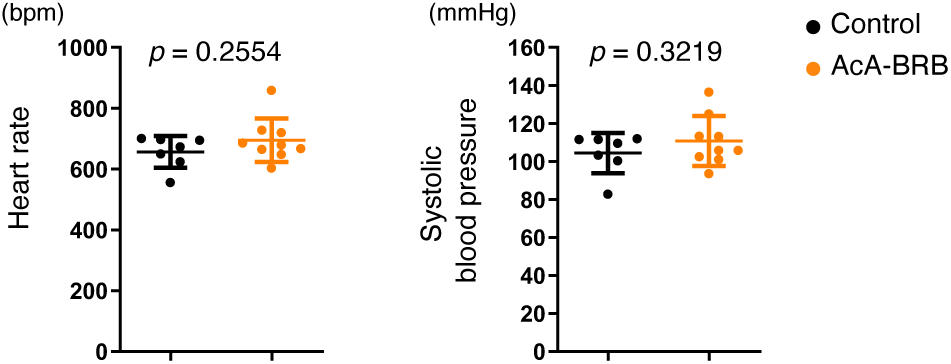
Heart rate and blood pressure in berberrubine-treated aged mice. Heart rate (bpm, left) and systolic blood pressure (mmHg, right) in 24-month-old WT mice after 3 months of treatment with berberrubine or tap water (control) (n = 7, control; n = 9, AcA-BRB). Statistical significance was determined using a two-sided unpaired Student’s *t*-test.

